# Global change factors reshape the links between litter properties, decomposers, and decomposition in mature oak forests

**DOI:** 10.1101/2025.07.18.665400

**Authors:** Matthew L. Meehan, Mathilde Chomel, Zhabiz Vilkiji, Katy J. Faulkner, Tancredi Caruso, A. Robert MacKenzie, Virginie Baldy, Richard D. Bardgett, David Johnson

## Abstract

1. Increasing atmospheric CO_2_ concentrations alongside more frequent and severe droughts are key global change factors impacting litter decomposition and global carbon cycles. However, we have a poor understanding of how both drought and elevated CO_2_ impact interactions between initial litter chemical properties and abiotic and biotic components of the decomposition environment, especially under field conditions.
2. We close this gap by testing how drought and elevated atmospheric CO_2_ concentrations modify rates of litter decomposition by acting through two pathways, the decomposition environment and initial litter properties. For this analysis, we used two long-term manipulative field experiments in mature oak woodlands: the Birmingham Institute of Forest Research Free-Air CO_2_ Enrichment (BIFoR FACE) experiment and the Oak Observatory at the Observatoire de Haute-Provence (O_3_HP) drought experiment that reduces seasonal precipitation. In each experiment, litter bags were deployed in a reciprocal transplant design whereby control and experimental litter (i.e. litter derived from plots under either drought or elevated CO_2_ conditions) were incubated in control and experimental plots at the respective experiment. We measured litter mass loss, functional groups, C:N ratios, moisture content, and microbial and mesofauna communities.
3. We found that drought and elevated CO_2_ concentrations modified mass loss rates through different pathways. In the drought experiment, litter placed in droughted conditions decomposed slower as the decomposition environment shaped mass loss rates more than initial litter properties. By contrast, both decomposition environment and initial litter properties impacted litter C:N, functional groups, and microbial communities during the experiment. In the elevated CO_2_ experiment, experimental litter derived from elevated CO_2_ plots corresponded with slower mass loss over the first three harvests as the decomposition environment did not impact litter properties or decomposers.
4. *Synthesis*: We found that drought and elevated atmospheric CO_2_ can modify mass loss rates through distinct pathways. Further research is needed to disentangle the relative roles of climate and decomposers with decomposition rates under drought and elevated CO_2_, but our experiment highlights that litter decomposition will be reshaped by global change factors through multiple mechanisms.

## Introduction

Forest ecosystems cover 31% of global land cover (FAO, 2020) and play a critical role in global carbon (C) cycling (Harris et al., 2021) as global forests take up 1.1 ± 0.8 Pg C year^-1^ (Pan et al., 2011). Yet these ecosystems are being reshaped by global change factors, including increasing atmospheric CO_2_ concentrations and increased frequency and intensity of extreme weather events such as droughts (IPCC, 2021). Elevated CO_2_ and drought independently affect C cycling because forests exhibit cellular- to ecosystem-level responses to both factors (Arain et al., 2022; Blaschke et al., 2001; Cech et al., 2003; Lee et al., 2021; Martínez-Sancho et al., 2022; Norby et al., 2024; Reay et al., 2025). For example, in a mature stand of *Eucalyptus tereticornis*, elevated CO_2_ increased C uptake (through gross primary production) by 12%, although net gains were lost due to an increase in soil fluxes, including from respiration (Jiang et al., 2020). In addition, a summer drought in 2022 resulted in nearly half of Europe’s soil moisture levels becoming critically low (Toreti et al., 2022), which corresponded with a reduced C uptake of ∼60 TgC (van der Woude et al., 2023). Because drought and elevated atmospheric CO_2_ concentrations affect ecosystem C exchange, it is critical to understand how these global change factors influence key ecosystem processes connected to C cycling, such as litter decomposition.

Litter decomposition is driven by soil biota, which break down litter into simplified compounds, leading to plant uptake of nutrients (Krishna & Mohan, 2017), the release of by-products into the atmosphere, and incorporation of nutrients into various soil pools (Lavallee et al., 2020). Litter biota include microbes, bacteria and fungi, that mineralize litter and organic matter *via* the secretion of enzymes (Romaní et al., 2006) and meso- and macrofauna who fragment litter but also graze on microbes (Coleman et al., 2018; Seastedt, 1984). Globally, litter decomposition is also driven by climatic factors, which regulate the activity of decomposers, such as temperature and annual precipitation, along with initial litter properties such as C:N ratio, lignin and tannins (Wu et al., 2025; Zhang et al., 2008). Both elevated CO_2_ and drought can impact litter decomposition through modifications to initial litter properties (hereafter, litter origin) and by modifying the environment (hereafter, the decomposition environment). However, the mechanisms, direction, and magnitude of responses will likely differ substantially between drought and elevated CO_2_.

Drought strongly modifies the belowground environment as reductions in moisture content have cascading effects on soil processes and microbial and animal communities. Here, desiccation has differential effects on particular groups of organisms: for example, fungi are generally more resistant to drought than bacteria (Canarini et al., 2024; de Vries et al., 2018; Yuste et al., 2011). Martin et al. (2024) found that reduced precipitation decreased soil animal abundance by 35% as species and group traits will dictate their tolerance to drier conditions (Lindo et al., 2012). The effects of elevated CO_2_ on decomposer communities are more subtle and often variable (Blankinship et al., 2011). Although not nearly as intense as drought, elevated CO_2_ concentrations can influence the soil microclimate, notably soil moisture (Leuzinger & Körner, 2007), which may impact decomposers. But a further distinction between elevated CO_2_ and drought on the decomposition environment is from changes in C allocation belowground. Elevated CO_2_ and drought have contrasting effects on the flux of recent plant photosynthate, which in forests is a major pathway of energy flow to soils (Högberg et al., 2001; Johnson & Griffiths, 2026). Under elevated CO_2_, trees increase belowground inputs of C: for example, at the Duke Forest FACE experiment, fine root biomass increased by 23%, and microbial biomass by 15% in response to sustained elevated CO_2_ (Drake et al., 2011). In a mature oak forest, similar increases in below-ground productivity was observed, including in the production and turnover of mycorrhizal mycelium (Reay et al., 2025). Increasing belowground C inputs may impact decomposition rates indirectly through altered biological activity, including soil enzymatic rates (Carney et al., 2007; Ochoa-Hueso et al., 2017) or changes to the ectomycorrhizal community, altering decomposition rates (Lindahl et al., 2021). By contrast, droughted trees tend to fix and allocate less C below-ground (Hagedorn et al., 2016), until their recovery where plants then allocate more C to roots to facilitate water and nutrient uptake (Gao et al., 2024; Joseph et al., 2020).

Both drought and elevated CO_2_ concentrations can affect leaf properties, which may subsequently impact the carbon and nitrogen content of senesced litter. Under drought, carbon and nitrogen content can shift through increased sugar accumulation and shifting primary and secondary plant metabolites (He et al., 2020; Mutanda & Mafirakurewa, 2026). Decreasing N concentrations in plants is a common response to enriched CO_2_ concentrations (Yuan & Chen, 2015) as N dilution within leaves together with C can alter the C:N ratio of the eventual litter (Ainsworth & Long, 2005; Taub & Wang, 2008). In addition, finer-scale litter chemical properties, such as phenol and polyphenol concentrations, captured by mass-spectrometry based metabolomics are also important as regulators of decomposition and impacted by these perturbations (Kasurinen et al., 2007; Quer et al., 2022). These and other changes to litter properties under both drought and elevated CO_2_ can affect litter mass loss rates (Cotrufo et al., 2005; Zhang et al., 2025) and decomposer communities (Allison et al., 2013; Martiny et al., 2017; Nielsen et al., 2024). Together, alterations to the decomposition environment and initial litter properties will impact litter decomposition. However, what remains difficult to assess is the relative importance of these two pathways in governing mass loss rates under drought and elevated CO_2_.

Disentangling the effects of initial litter properties from those of the decomposition environment loss under field conditions is challenging but doing so better captures interactions between these two pathways on litter decomposition. For instance, Prieto et al. (2019) found greater reductions in decomposition when experimental-derived litter (i.e., warming and reduced rainfall) decomposed in treated plots, relative to the control litter. Here, interactions between decomposition environment and litter origin led to slower decomposition than could be predicted with litter origin or decomposition environment alone. Overall, both drought and elevated atmospheric CO_2_ have the potential to affect litter decomposition *via* alteration of litter properties and the decomposition environment (which includes decomposers and litter moisture content) but how initial changes to litter properties interact with the decomposition alongside changes to the decomposer community *in situ* remains poorly understood. Resolving this uncertainty requires large-scale experimental infrastructure with long-term history of treatments in combination with multi-variate analysis of litter properties and other factors.

We investigated how drought and elevated atmospheric CO_2_ independently affect litter decomposition in mature forests by conducting reciprocal transplants within two, large-scale manipulative field experiments: the Birmingham Institute of Forest Research Free-Air CO_2_ Enrichment (BIFoR FACE) experiment and the Oak Observatory at the Observatoire de Haute-Provence (O_3_HP) drought experiment. We propose that drought and elevated CO influences litter decomposition through two distinct pathways. The first pathway involves changes to the decomposition environment and involves testing decomposition of the same litter in contrasting environments (i.e. control vs. drought and control versus elevated CO_2_ plots). These effects capture how changes in soil moisture, microclimate, and decomposer communities directly regulate decomposition. The second pathway involves litter origin, which occurs when different treatments (e.g. ambient rainfall versus drought, or ambient CO_2_ versus elevated CO_2_) change litter properties prior to decomposition, but which carry-over to affect the decomposition process. For example, plant growth under drought and elevated CO_2_ conditions may modify litter properties prior to litterfall and subsequent decomposition. Specifically, we focus on impacts on C:N ratio, and a range of metabolites because these properties have been demonstrated to regulate decomposition in past work. Within the field experiments, we aimed to uncover the relative strength of each pathway that governs mass loss rates. We hypothesized that drought and elevated CO_2_ would modify mass loss rates in our study with the relative strength of the two pathways varying between the global change factors. We predicted that amplified drought conditions would alter mass loss more through the decomposition environment due to the greater influence of drought on the soil microclimate and decomposer community. Conversely, we predicted that elevated CO_2_ concentrations would influence mass loss relatively more through long-term (carry over) effects given the weaker links between the decomposition environment and mass loss and but still present direct effects of elevated CO concentrations on initial litter properties. Our reciprocal transplant design at BIFoR FACE and O_3_HP allowed us to test the relative strength of these two pathways and whether interactive effects between litter origin and decomposition environment influence decomposition rates when subject to elevated CO_2_ and drought.

## Methods

### Overview

Within each experimental facility, we collected control and experimental litter and in January 2020 transplanted the two litter types into control and treated plots (Supplementary Information Figure S1). We harvested litterbags across four time points and measured (across all or a subset of time points) litter mass loss, litter functional groups, properties and metabolomics, and the microbial (phospholipid fatty acid analysis; PLFA) and animal community structure (the abundance of mites, collembolans and other larger macrofauna) directly associated with the recovered litter. Note that we did not transplant litter between the two experiments (i.e., litter from BIFoR FACE was not transplanted at O_3_HP and *vice versa*). We only transplanted litter within each experimental facility between the control and experimental apparatus, but we will continue to refer to it as a reciprocal transplant experiment.

### Experimental sites

The reciprocal transplant study used the BIFoR FACE (52.801°N, 2.301°W) elevated CO_2_ experiment in the United Kingdom and the O_3_HP (43.936°N, 5.719°E) drought experiment in southern France (Figure S1). Both are unique, long-term experiments comprising large-scale, costly infrastructure. The BIFoR FACE facility started in 2017 on a *Quercus robur* L. forest that was planted around 1850 (Hart et al., 2020; MacKenzie et al., 2021) containing coppice *Corylus avellana* L. and abundant self-set *Acer pseudoplatanus* L. understory. The CO_2_ treatment began in April 2017 and comprises six infrastructure plots (elevated CO_2_, n = 3, ambient CO_2_, n = 3). Each plot is approximately 30 m in diameter and comprises 16 peripheral towers and one central tower that extend ∼ 1m above the canopy. In the elevated CO_2_ plots, an additional 150 ppm of CO_2_ above ambient (i.e. ∼36 % increase over ambient) is emitted laterally from the ground to the forest canopy, while ambient CO_2_ arrays mimic the treatment but only receive ambient air. The increase of CO_2_ concentrations begins at the soil and across the entire vertical profile of the plot (Hart et al., 2020), meaning litterbags would be incubated in an enriched CO_2_ environment. The towers release CO_2_ from ∼April 1 to October 31 each year, coinciding with oak bud burst and last leaf fall (Figure S2). The +150 ppm of CO_2_ is similar to other large-scale experimental facilities such as EucFACE (e.g., Jiang et al., 2020) and represents the projected mid-century CO_2_ concentration, regardless of model scenario (Friedlingstein et al., 2014). It also allows for ecophysiological effects to be detected with both aboveground and belowground measurements (Gardner et al., 2022; Norby et al., 2024; Reay et al., 2025) and provides an end-member of the small set of forest FACE treatments (Norby, 2025). For this experiment, when we refer to plot in the statistical analysis, we refer to the six arrays where litterbags were transplanted.

The O_3_HP drought experiment started in 2012 on *Quercus pubescens* Willd forest, containing *Acer monspessulanum* L. (75% vs. 25% coverage, respectively) and *Cotinus coggygria* Scop understory. A rain exclusion device, measuring 15 m x 20 m excludes ∼30% of precipitation at canopy top, relative to the nearby control plot. In our study, the rain exclusion device was operational between May to September and the total precipitation was 40% lower in rain exclusion plots over the course of the experiment (total precipitation, control: 1652 mm, drought: 997 mm; Figure S3) (Aupic-Samain et al., 2021; Gauquelin et al., 2011; Santonja et al., 2017). There is only a single drought apparatus established at O_3_HP although the scale of the apparatus brings many benefits (Davies & Gray, 2015) and enables creation of ‘plot-like’ conditions relevant to the scale of litterbags and the key processes governing their decomposition. The bags were therefore placed in a clumped distribution under and outside of the rain exclusion devices (minimum 4m apart) to create three experimental plots within the treatment. This helped capture how changes in the microclimate within segregated areas *via* amplified drought can impact litter mass loss, litter properties, and decomposers within individual litterbags. When we refer to plot in the statistical analysis for this experiment, we are referring to these clumped patches of litterbags. The use of equivalent experimental designs at both O_3_HP and BIFoR FACE means that the effects of decomposition environment and litter origin on litter decomposition in response to drought and elevated CO_2_ can be tested and compared with the maximum possible rigour given the constraints imposed by the disparate locations of the experimental platforms.

### Litterbag preparation and sampling

*Quercus* litter was sampled from spatially separated areas of each site in BIFoR FACE and O_3_HP in October 2019 and shipped in coolers to The University of Manchester and air dried. We measured the %C, %N, C:N and metabolomic signatures on the initial litter (Quer et al., 2022; see *Metabolomic signature of initial litter* in Methods S1 for more details). Litterbags comprised of 1.8 mm nylon mesh with two separate compartments (Supplementary Information Image S1), each receiving 3 g of dried litter. One compartment was used for chemical analysis of litter and assessment of microbial community composition, decomposition, and moisture content. The other compartment was used solely to characterize soil fauna communities. A total of 480 litterbags were used (i.e. 240 litterbags for BIFoR FACE and 240 for O_3_HP). For each experimental site, we had 2 decomposition environments (control and drought or elevated CO_2_) x 2 litter origins (control and experimental) x 3 plots x 4-time points x 5 replicates = 240). The experiment began January 2020 where litterbags were placed on the experimental sites. Specifically, litterbags were placed in pairs consisting of one with experimental litter, and the other with control litter. Freshly fallen litter was removed prior to placing the litterbags on the ground surface and then replaced over the litterbags. Litterbags were then fixed with two galvanized nails to prevent movement by animals or wind. Litterbags were harvested across four time points at both experimental sites. At BIFoR FACE, litterbags were harvested September 2020 (8 months; Harvest one; H1), January 2021 (12 months; Harvest two; H2), June 2021 (17 months; Harvest three, H3) and December 2021 (23 months; Harvest four, H4), corresponding to the fourth and fifth year of activation of the elevated CO_2_ treatment. Four litter bags could not be retrieved from BIFoR FACE at H4 due to a tree limb falling and disturbing litter bags. At O_3_HP, litterbags were harvested October 2020 (10 months; H1), January 2021 (13 months; H2), September 2021 (21 months; H3) and October 2022 (34 months; H4). The duration of decomposition differed between experimental sites due to variation in decomposition rate. We initially aimed for ∼24 months of decomposition but incubation in the drought experiment was extended by an additional year to achieve 50% mass loss.

At each sampling time, litterbags were shipped to The University of Manchester for processing. Because the litterbags were placed on the forest floor, at times we needed to gently remove extraneous debris that attached to litter during transit or while retrieving them. This was done solely for the remaining litter from the ‘chemical analysis compartment. The fresh weight (g) of the litter was taken before litter was frozen at −20 °C overnight and 48 hours of freeze-drying for determination of dry weight. For soil fauna, litter was removed from the compartment and placed on Tullgren funnels to extract individual animals for seven days into absolute ethanol. After extraction, litter was placed into ovens at 65 °C for two days before weighing to measure litter dry weight. Apart from litter mass loss rates (measured at all dates), litter properties and decomposer communities were measured from H3 onward, when ∼20-60% of the original litter remained, to capture drought and elevated CO_2_ effects during mid to late stage of decomposition.

### Litter decomposition, properties, chemistry, and moisture content

Litter decomposition was calculated as litter mass loss using the following equation:

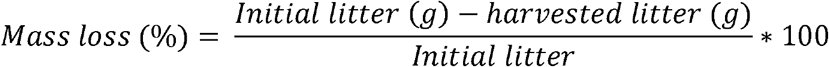

We also calculated the exponential decomposition rate coefficient (*k_t_*) at each harvest point using the equation from (Olson, 1963):

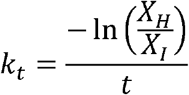

Where *X_H_* is the weight (in g) of harvested litter, *X_I_* is the weight of initial litter (in g), and *t* is time in years since the beginning of the experiment. Litter decomposition and litter moisture was measured from all litterbags across all four time points. We measured the %C, %N and C:N ratio of initial and decomposed litter with an Elementar Vario EL element analyser (Hanau, Germany) on 5 mg of ball-milled litter (Retsch Ball Mill MM 400, Haan, Germany) at both sites.

We also measured litter chemistry (specifically, characterizing C-containing functional groups) of H3 and H4 litter from both sites using Fourier transform infrared (FTIR) spectroscopy (Bell et al., 2018). FTIR can reveal changes to litter functional groups associated with cellulose, hemicellulose and lignin and indicate loss and transformation of these compounds during decomposition (Lammers et al., 2009; Liu et al., 2016; Turrión et al., 2025), and thus has potential to detect litter chemical properties and pathways that are missed with C and N alone. Ball-milled litter samples were placed onto a Universal Attenuated Total Reflection (UATR) spectrometer diamond crystal to cover the 2-mm window, and the absorbance (A cm^-1^) was measured within a 650 to 4000 A cm^-1^ range, at 4 cm^-1^ resolution and 16 scans per sample. We also ran a background correction after every ten samples. We applied a baseline correction for each spectra (i.e., sample) using the modpolyfit method in the R package ‘baseline’. To standardize values across our samples, we Z-transformed each spectra, where values were transformed to a mean of 0 and SD of 1 by subtracting by its own mean and dividing by its own standard deviation (akin to a Standard Normal Variate normalization) to reduce multiplicative scatter effects (Bell et al., 2018; Jardine et al., 2017). We then calculated the area of the curve for each respective functional group using a shoelace formula (Wang et al., 2023). The assignment of wave number regions to leaf litter functional groups was based on Liu et al., (2016) who used FTIR to examine functional groups across four deciduous litter origins, including *Quercus robur* (Supplementary Information Table S1).

### Phospholipid fatty acid analysis (PLFA)

We used PLFAs to measure how fungal and bacteria concentrations and the general microbial community were impacted by drought and elevated CO_2_ in leaf litter at both sites (Buyer & Sasser, 2012). Due to high litter mass loss rates in the fourth harvest of BIFoR FACE, we did not have enough material remaining to quantify microbial communities through PLFAs. So, we elected to measure PLFAs for both sites only on litterbags collected in the third harvest. We quantified PLFAs with ∼0.4-0.5 g of ball milled, freeze-dried litter. Four ml of Bligh-Dyer extractant containing C19:0 of an internal standard was added to each sample to extract lipids. Once extracted, we used a gas chromatograph (Agilent 7890A Gas chromatograph, Santa Clara, CA, USA) to assign and calculate peak height, where PLFA group concentrations were expressed as nmol g dry weight of litter^-1^. We used 18 fatty acid markers to characterize microbial communities (Frostegård et al., 1993; Hill et al., 2000) on leaf litter, consisting of 6 and 4 markers for gram positive (i14:0, i15:0, a15:0, i16:0, i17:0, a17:0) and gram negative (16:1ω7c, cy17:0, 18:1ω7, and cy19:0) bacteria, respectively. In addition, we used 15:0 and 17:0 as general bacteria markers, and 10Me16:0, 10Me17:0, and 10Me18:0 as markers for actinobacteria. The marker 18:2ω6,9 was the sole fungal marker in our experiments. Fatty acids 14:0 and 16:0 were used as general microbial markers in our study, not specific to fungi or bacteria. With these markers, we analyzed decomposition environment and litter origin effects on total PLFA concentrations (proxy for microbial biomass), F:B ratio (by dividing 18:2ω6,9 abundance by the summed total of bacteria markers) and PLFA community composition across both experiments.

### Soil fauna community

Soil fauna was sorted and counted into higher taxonomic or mixed groups, comprising: Collembola, Oribatida, Astigmata, Prostigmata, Mesostigmata, and ‘other’ fauna, which included Annelida, Isopoda, and Insecta. Soil fauna abundances were standardized by the dry weight of litter (individuals g dwt^-1^), to account for differences in sample mass prior to analysis. Although found in litterbags, macrofauna counted for less than 5% (on average) of total abundances in our study, and as a result most individuals captured were mesofauna, particularly mites and collembolans.

### Statistical analysis

All statistical analyses were performed with the program R (version 4.3.1) (R Core Team, 2023) and figures were produced with the package ‘ggplot2’ (Wickham, 2016). To analyze for differences in initial litter properties between litter types (litter from control and experimental plots) prior to the experiment beginning, we used one-way Analysis of Variance (ANOVA) tests and Permutation Multivariate ANOVAs (PERMANOVAs) to examine for differences between C%, N% and C:N ratio (ANOVA) and leaf metabolomic profile (PERMANOVA, with Euclidean distance) between our treatments (ambient vs. drought or elevated CO_2_). To do this, we used the aov function in base R and the adonis2 function in the package ‘vegan’ (Oksanen et al., 2022), respectively. Here, we partitioned the data by experiment (BIFoR FACE and O_3_HP) and ran separate models for both.

We examined the effect of decomposition environment (ambient vs. drought or elevated CO_2_) and litter origin (control vs. experimental) on litter mass using linear mixed models where we partitioned the data between the two experiments (BIFoR FACE and O_3_HP). To do this, we ran a linear mixed model with decomposition environment (ambient vs. drought or elevated CO_2_), litter origin (control vs. experimental), harvest, and their interactions as the fixed effects, and experimental plot as the random effect. After this, we adapted the approach from Nielsen et al. (2024) to gauge the strength of both decomposition environment and litter origin at each time point, where we parameterized the effect of each treatment using log response ratios stemming from the mean response of model predictions. We also calculated standard errors from model predictions for each log response ratio using the formula provided in Hedges et al. (1999) where log response ratios were considered significant when the confidence interval (mean ± 2 standard errors) did not cross zero. To calculate effect sizes and confidence intervals, we adapted from Powell, (2024) where we used the function ggemmeans from the package ‘ggeffects’ (Lüdecke, 2018) to extract model predictions and the ‘pairs’ and ‘emmeans’ functions (from the package emmeans) (Lenth, 2025) to generate p-values associated with each effect size.

For the remaining univariate models, we analyzed experiments (BIFoR FACE and O_3_HP) and time points separately, in part, due to diverging litter and microbial properties between the two experimental locations (Figure S4). We used linear mixed models where the fixed factors were decomposition environment, litter origin, and their interaction, and experimental plots as a random factor. We used linear mixed models for %C, %N, C:N ratio, total PLFA concentration (nmol g dwt^-1^), fungi : bacteria ratio, animal total and group abundances (individuals g dwt^-1^), which includes Collembola, Oribatida, Astigmata, Mesostigmata, Prostigmata, and ‘other’ fauna. Note, we did not analyze Astigmata abundances in the drought experiment as low abundances were present in samples. For all linear mixed models described above, we used the lmer function in the package ‘lme4’ (Bates et al., 2015) where we calculated F-statistics and p-values with Satterthwaite’s method using the package ‘lmerTest’ (Kuznetsova et al., 2017). If model residuals were not normally distributed, we ln or arscine-transformed the dependent variable and re-ran the model. Occasionally, we found singularity within our linear mixed models as the variance within our random factor terms was extremely low (essentially zero) and the model was overfit. When this occurred, we removed the random factor, and we re-ran the model without the random term. We found this had no effect on p-values (because experimental plot explained little variance within the linear mixed model), so we reported the ANOVA results. We provided marginal and conditional R^2^ for linear mixed models (using the function r.sqauredGLMM from the package “MuMIn” (Barton, 2025)), and adjusted R^2^ for linear models, when the random factor was removed.

For multivariate variables, including leaf litter functional groups (FTIR) and microbial community composition, we ran PERMANOVAs using the adonis2 function in the package ‘vegan’ (Oksanen et al., 2022). We ran PERMANOVAs at both the litterbag-level (n = 56 or 60), where individual litterbags were the values, and the subplot-level (n = 12), where averages values were calculated for litterbags of the same litter origin, within each plot. Because of the limitations of performing multivariate statistics with our experimental design, there is inherent pseudo-replication within our analyses. But by conducting tests at both the litterbag and subplot-level, we were able to measure the impact of decomposition environment and litter origin across two scales to determine the impact of both treatments on litter-C functional groups and microbial community composition. For leaf litter functional groups, we used Euclidean distances within our models, where the functional groups were z-transformed (mean = 0, sd = 1) to standardize them before calculating distances. With microbial communities we used Bray-Curtis dissimilarity as it better suited for compositional data (like the abundances of each PLFA peak). Each model was fitted with the same fixed factors, consisting of decomposition environment, litter origin and their interaction. If the interaction was not significant within these models, we removed the interaction and re-ran the model. To visualize leaf litter functional groups and microbial communities we used a Principal Component Analysis (PCA) ordination plot using the rda function in the package ‘vegan’ (Oksanen et al., 2022). For microbial communities, we Hellinger-transformed datasets using the decostand function in the ‘vegan’ package prior to the PCA.

## Results

### Initial litter properties and metabolomic profiles

We found strong differences between the C:N ratio of the initial litter collected at control and droughted plots at O_3_HP (F_1,8_ = 13.92, p = 0.006; Table S2), with the C:N ratio of control litter exceeding that of the droughted litter. Furthermore, strong differences were detected between the metabolomic profiles between the initial control and droughted litter (R_2_ = 0.25, F_1,8_ = 2.62, p = 0.009; Figure S5). Control litter from the drought experiment was mainly composed of ellagitannins (from C_34_H_24_O_22_ to C_56_H_40_O_31_), catechin (C_15_H_14_O_6_), and ellagic hexoside (C_20_H_16_O_13_) that were overexpressed within litter. Other metabolites such as quercetin hexosides (or rhamnoside), kaempferol hexosides, and isorhamnetin hexosides were at similar quantities between control and drought treatment. By contrast, there were no differences in C:N ratio (F_1,8_ = 0.001, p = 0.977; Table S2) or metabolomic profiles (R_2_ = 0.14 F_1,8_ = 1.34, p = 0.116; Figure S5) between litter collected from the ambient and elevated CO_2_ plots at BIFoR FACE.

### Elevated CO_2_ experiment at BIFoR FACE

We found that during the first eighteen months of the decomposition experiment, litter from elevated CO_2_ plots (i.e., experimental litter) decomposed slower than litter from control plots (litter origin: F_1,215.09_ = 5.93, p = 0.016) through H1-3 (time: F_3,215.47_ = 155.97, p < 0.001) until H4 in both decomposition environments (Figure 1A, Table S3 contains exponential decomposition rate coefficient, *k_t_*, values). The effect of litter origin was consistent albeit weak as effect sizes did not differ from zero for all harvests (Figure 1B), matching p-values (p > 0.10). Mass loss rates slightly increased for litter bags in the elevated CO_2_ environment at the third harvest (Figure 1C), which differed from the remaining time points, although this was not significant for both the control (p = 0.08) and experimental (p = 0.22) litter.

**Figure 1.**
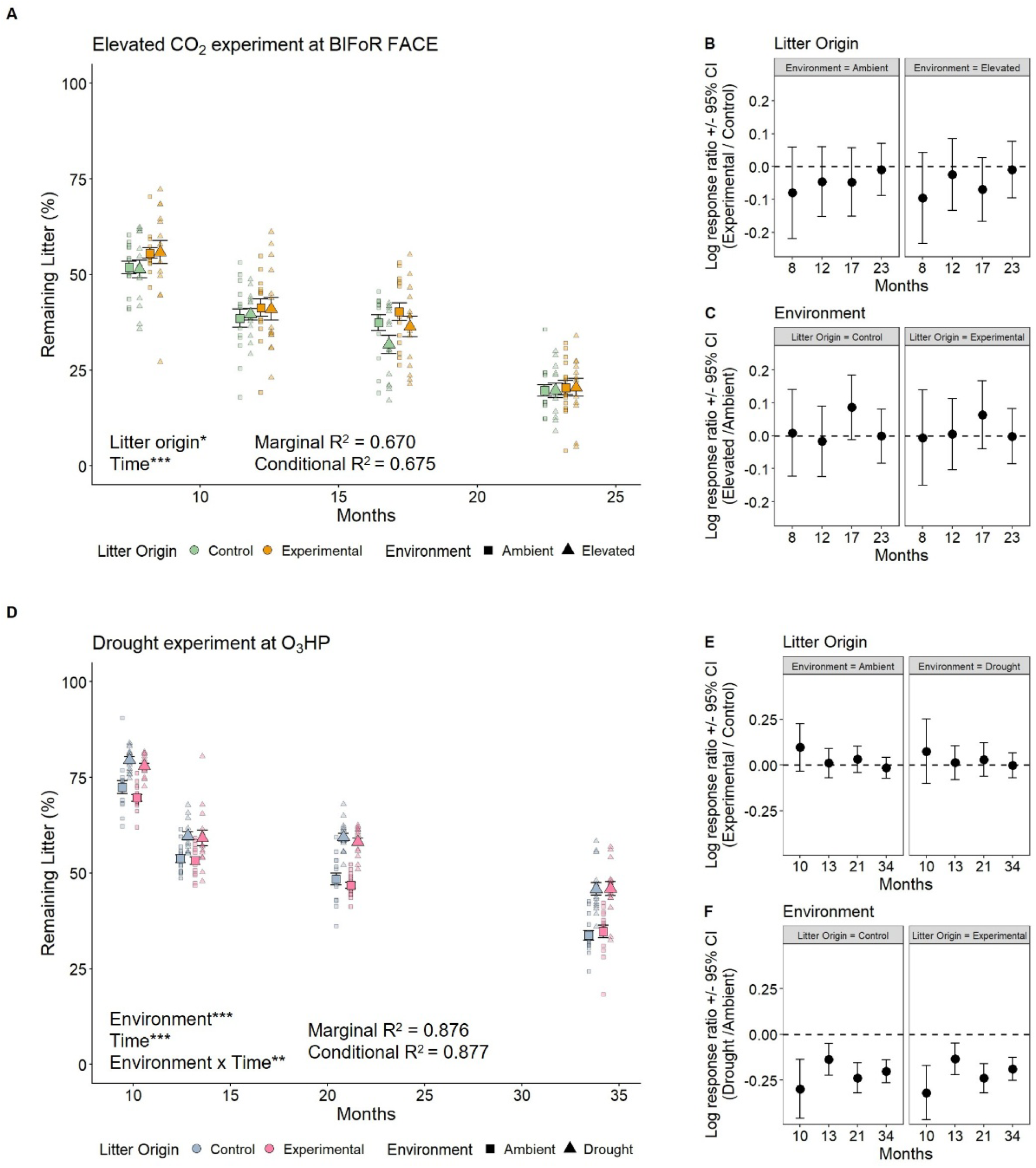
Litter decomposition (shown as % remaining litter) for the elevated CO_2_ experiment (A) and drought experiment (D). The x-axis indicates the number of months since initiation of the experiment and differs for upper and lower panels. Larger points represent mean (± SEM) while smaller points are the raw data values for individual litterbags. * = p < 0.05, ** = p < 0.01, and *** = p < 0.001. We have only provided p-values of factors that were statistically significant. Litter bags in the elevated CO_2_ experiment were harvested after 8 (H1), 12 (H2), 17 (H3) and 23 (H4) months, while litter bags in the drought experiment were harvested after 10 (H1), 13 (H2), 21 (H3), and 34 (H4) months. To improve plot interpretation, points were positioned with greater spacing on the x-axis. Log response ratios (± 2 standard errors) for the elevated CO_2_ experiment (B and C) and the drought experiment (E and F) are provided for each treatment, litter origin (B and E) and decomposition environment (C and F). The values for the response ratios have been partitioned by the remaining treatment. E.g. Response ratio for decomposition environment is shown for both control and experiment litter. Positive values indicate decomposition is greater for experimental litter or in treated plots (amplified drought or elevated CO_2_ concentrations). Negative values indicate the opposite. Response ratios were considered significant if error bars did not cross zero.

The C:N of litter was slightly higher in experimental litter compared to controls (F_1,56_ = 6.03, p = 0.02) at H3, but not at H4 (Table S4 and S5). This finding was partly reflected by FTIR spectroscopy analysis that showed that litter functional groups varied weakly between the litter origins at H3 at the subplot-level (R_2_ = 0.16, F_1,9_ = 1.87, p = 0.06; Figure 2A) but less so at the litterbag-level (R_2_ = 0.03, F_1,57_ = 1.84, p = 0.13; Figure S6A). At H4, we continued to find weak effects of litter origin on litter functional groups at the subplot-level (R_2_ = 0.16, F_1,9_ = 1.84, p = 0.09; Figure 2B) but found a slight difference between the litter types at the litterbag-level (R_2_ = 0.04, F_1,53_ = 2,10, p = 0.05; Figure S6B). Broadly, these results indicate a weak but present effect of litter origin on litter C functional groups. Neither decomposition environment nor litter origin affected litter moisture content at either H3 or H4 (Table S4 and S5).

**Figure 2.**
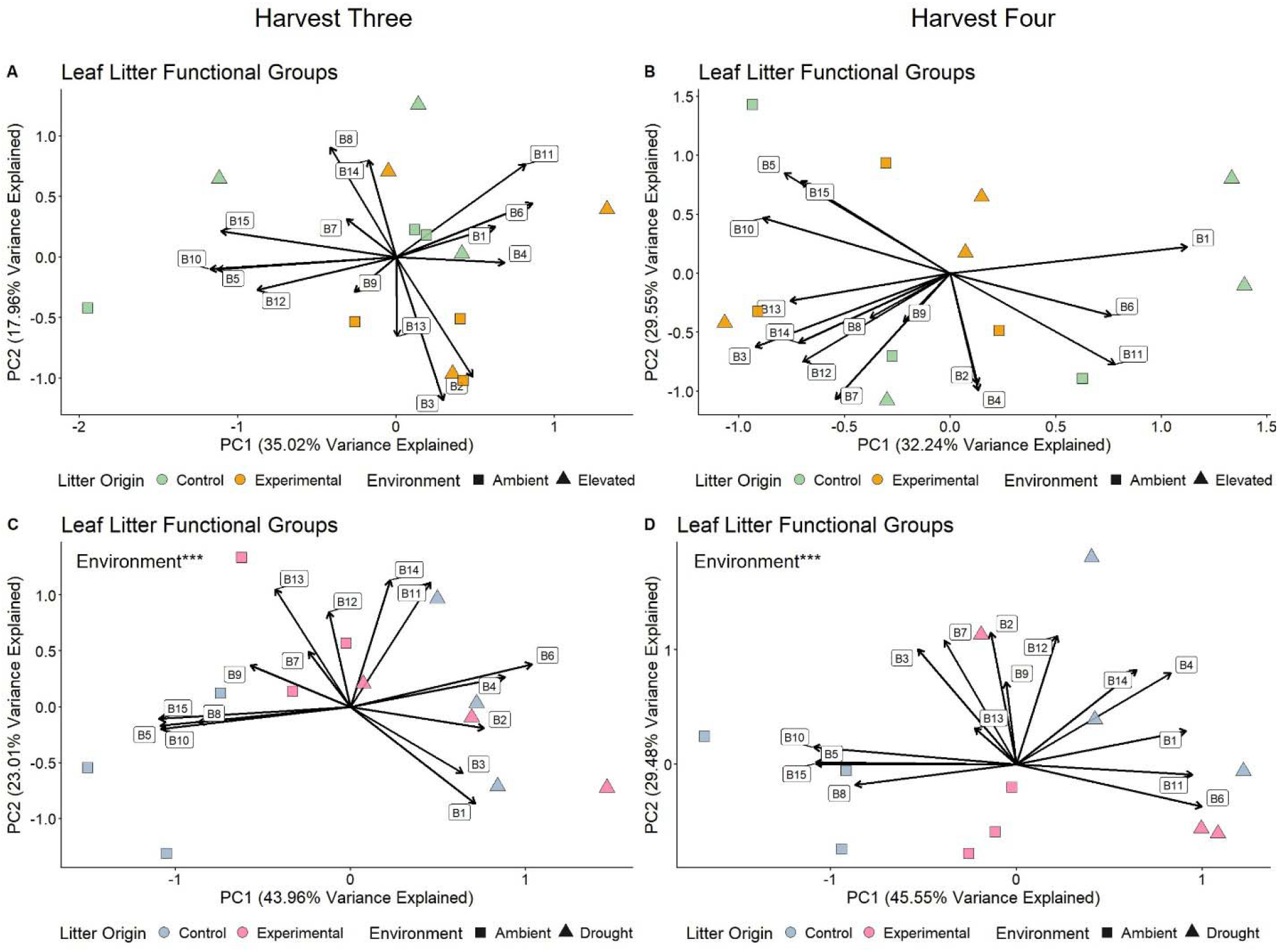
Principal component analysis (PCA) ordinations of leaf litter functional groups measured by Fourier transform infrared (FTIR) spectroscopy for the elevated CO_2_ experiment (top) and drought experiment (bottom) at harvest three (A and C, left) and four (B and D, right). Points represent average functional group values for treatments within a given plot, akin to the subplot-level. Corresponding functional groups for band numbers (B1, B2, B3, etc.) can be found in Table S1. *** = p < 0.001. We have only provided p-values of factors that were statistically significant.

Decomposition environment and litter origin did not influence PLFA abundance, the fungi : bacteria (F:B) ratio, and microbial community composition at H3 (Table S4, Figure S7). Likewise, no individual or total animal group abundances were affected by decomposition environment or litter origin in H3. In H4, however, the abundance of Mesostigmata (F_1,52_ = 5.04, p = 0.03) and Prostigmata (F_1,52_ = 7.30, p = 0.009) was significantly greater in control litter relative to experimental litter in both decomposition environments (Table S5).

### Drought experiment at O_3_HP

By contrast to elevated CO_2_, there were stronger effects of both decomposition environment and litter origin on measured parameters in response to drought. First, litter decomposition was slower in droughted plots relative to controls (decomposition environment: F_1,4_ = 147.91, p < 0.001) with remaining litter mass decreasing at varying rates (decomposition environment x time: F_1,220_ = 4.56, p = 0.004; Figure 1D; Table S3). This finding matches effect sizes as across all four harvests and two litter origins, decomposition was slower in the droughted decomposition environment (Figure 1F). Conversely, litter origin did not impact litter mass loss rates at any time point during the experiment (Figure 1E)

Remaining litter had lower C:N ratios in control plots (H3: F_1,4_ = 21.92, p = 0.009; H4: F_1,56_ = 37.85, p < 0.001) compared to drought plots. In addition, experimental litter (i.e., litter derived from drought plots) had lower C:N ratio, relative to the control litter in both decomposition environments (H3: F_1,52_ = 32.53, p < 0.001; H4: F_1,56_ = 9.54, p = 0.003) suggesting both decomposition environment and litter origin affected litter properties during the experiment (C:N but also %C and %N; Figure 3; Table S6 and S7).

**Figure 3.**
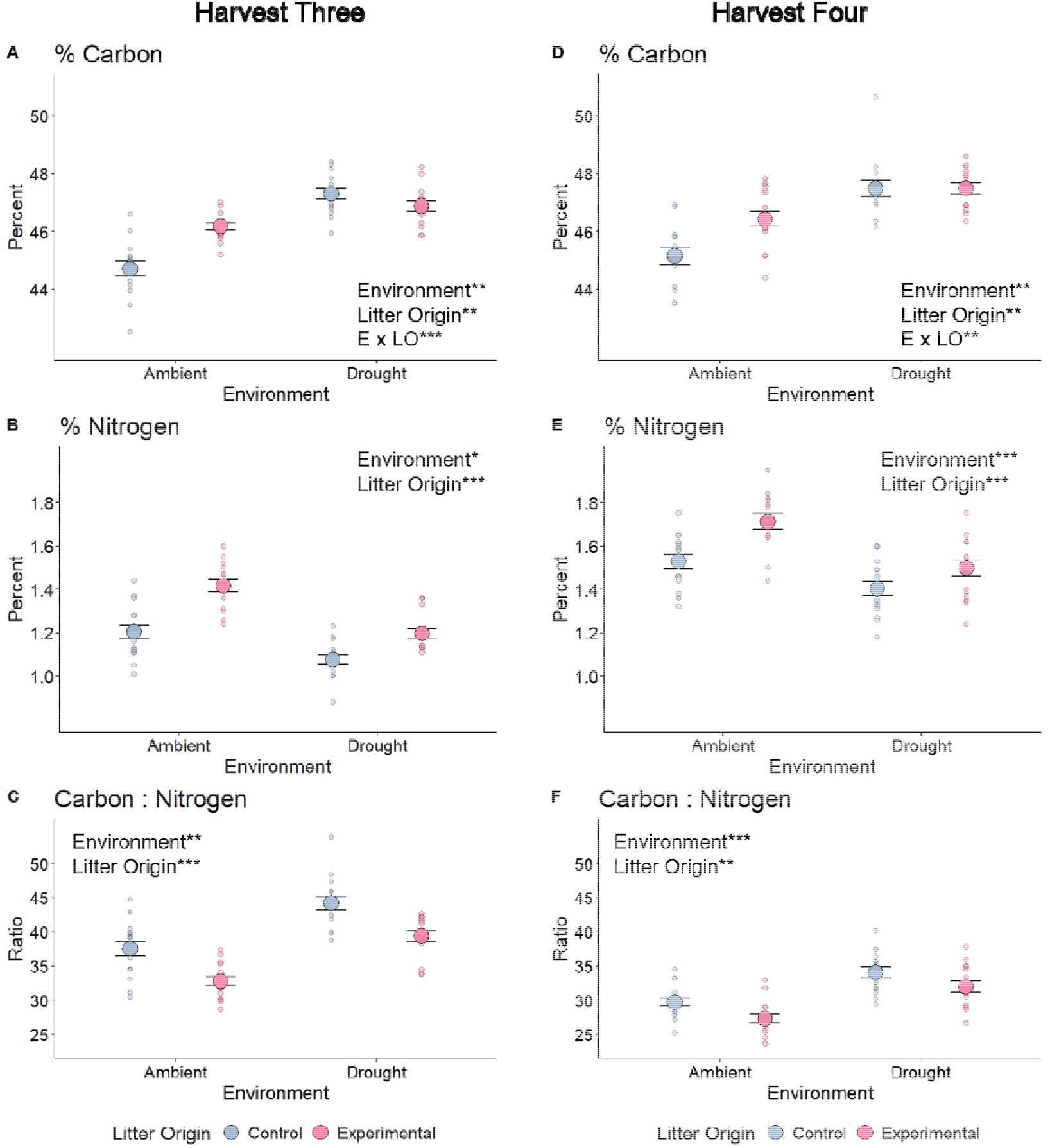
% Carbon (%C; A and D), % Nitrogen (%N; B and E) and C:N (C and F) of drought experiment litter sampled at the at third (left) and fourth harvest (right). Larger points represent mean (± SEM) while smaller points are the raw data values of individual litterbags. * = p < 0.05, ** = p < 0.01, and *** = p < 0.001. We have only provided p-values of factors that were statistically significant. E = Environment and LO = litter origin.

We observed a strong effect of decomposition environment on leaf litter functional groups. At the subplot-level, we found strong effects of decomposition environment at both H3 (R_2_ = 0.33 F_1,9_ = 5.48, p < 0.001) and H4 (R_2_ = 0.34. F_1,9_ = 5.48, p < 0.001) as reduced rainfall greatly impacted litter functional groups (Figure 2C-D). Within individual litter bags, we found strong interactions between decomposition and litter origin at both time points (H3: R_2_ = 0.04, F_1,56_ = 3.15, p = 0.006; H4: R_2_ = 0.03, F_1,56_ = 2.37, p = 0.04), indicating a weaker, but present effect of litter origin, alongside decomposition environment, on litter functional groups (Figure S6). We found divergent effects of the experimental drought on litter moisture content at O_3_HP. At H3, litter moisture content significantly decreased (by around 35%) in drought plots (F_1,4_ = 85.96, p < 0.001; Table S6). However, at H4, when litter moisture content in control plots was already low, there was no significant difference between the control and drought plots (F_1,56_ = 1.086, p = 0.30; Table S7).

After 21 months of decomposition (H3), experimental litter (i.e., litter from the droughted plots) had greater concentrations of PLFAs in both decomposition environment (F_1,52_ = 9.06, p = 0.004, Figure 4A), but the drought treatment reduced total PLFA concentrations (F_1,4_ = 15.69, p = 0.02) in litter regardless of its origin. The F:B ratio was greater in litter bags from the drought environment across both litter types (F_1,4_ = 19.42, p = 0.01, Figure 4B). At the subplot-level, decomposition environment strongly shaped microbial communities (R_2_ = 0.53, F_1,9_ = 14.82, p = 0.002, Figure 4C). Litter origin expressed a weak effect on PLFA composition (R_2_ = 0.15, F_1,9_ = 4.03, p = 0.07) at the subplot-level. But both litter origin (R_2_ = 0.08, F_1,57_ = 6.76, p = 0.004) and decomposition environment (R_2_ = 0.27, F_1,57_ = 23.99 p < 0.001) shifted PLFA composition at the litterbag-level (Figure 4D). Here, microbial composition in response to decomposition environments was largely explained by PC1 whereas microbial composition in response to litter origin was explained more by PC2. These axes reflected distinct PLFAs, with the fungal-related PLFA C18:2ω6,9 driving separation in both PC1 and PC2 (Figure 4D) with general bacteria and microbial markers (14:0, 15:0, 16:0 and 17:0) reflected litter origin differences on PC2. In addition, animal communities responded more to the decomposition environment than litter origin (Table S6 and S7). Mesostigmatid (H3: F_1,56_ = 31.18, p < 0.001) and prostigmatid mite (H4: F_1,56_ = 7.65, p = 0.007) abundances significantly decreased, and there was a trend for drought to reduce total animal abundances at H4 (F_1,4_ = 6.04, p = 0.07). Notably, there was a significant interaction of decomposition environment and litter origin on ‘other’ macrofauna abundances at H3 (decomposition environment x litter origin: F_1,52_ = 6.51, p = 0.01) where abundances were greatest in the droughted plots with control litter.

**Figure 4.**
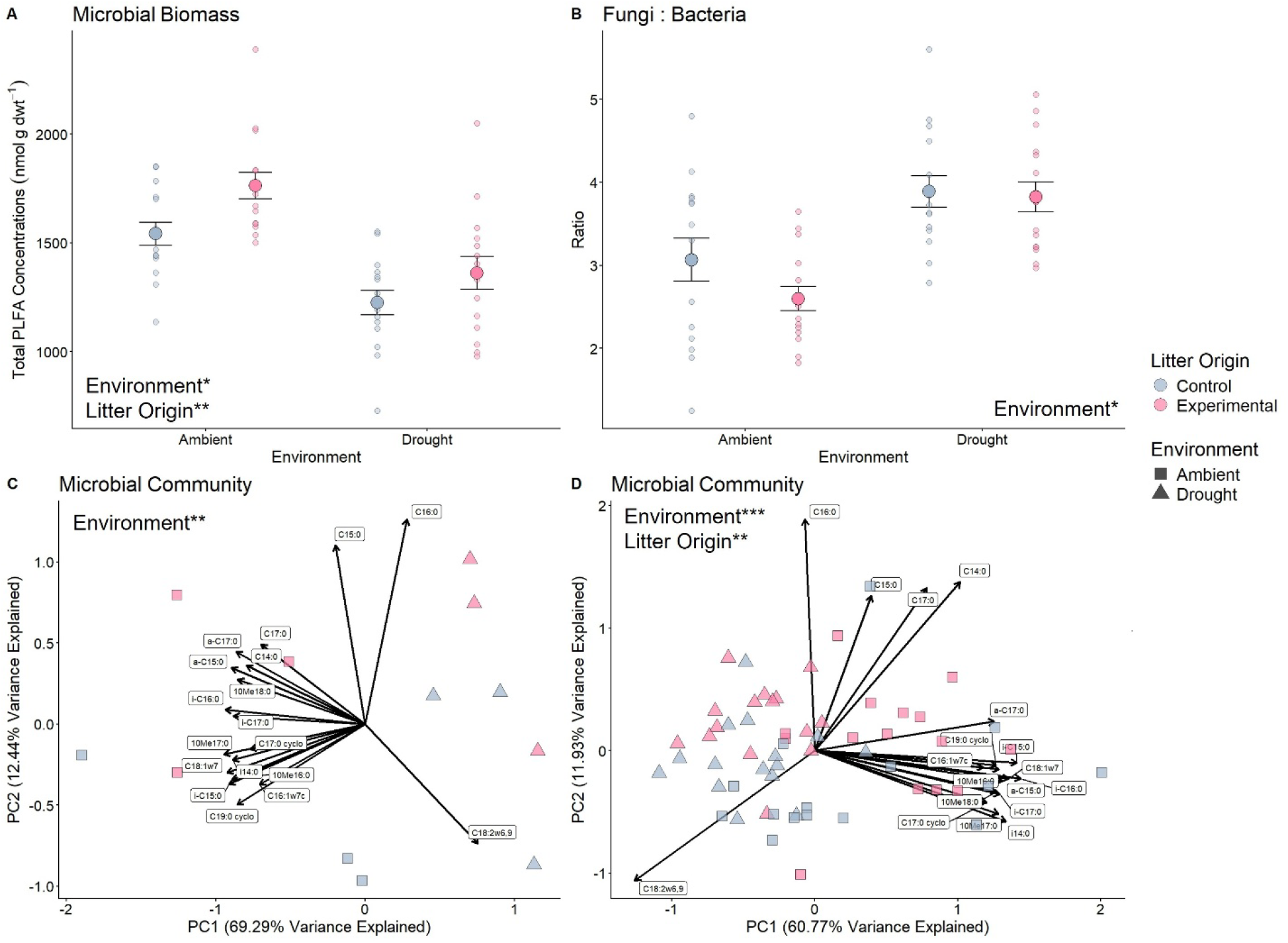
Microbial community indices as measured by phospholipid fatty acid analysis (PLFA) in litter for the drought experiment after 21 months (harvest 3). For A) (total PLFA) and B) (fungi : bacteria) larger points represent mean (± SE) while smaller points are the raw data values for individual litter bags. Principal component analysis (PCA) ordination of microbial community structure at subplot-level (C) and litterbag-level (D). * = p < 0.05, ** = p < 0.01, and *** = p < 0.001. We have only provided p-values of factors that were statistically significant.

## Discussion

The use of two large scale, long-term experimental manipulations of drought and elevated atmospheric CO_2_ provide results that address the need for ‘ecologically relevant’ (Rineau et al., 2019) investigations of the impact global change factors on soil C cycling processes. Our results reveal that long-term *in situ* manipulations of drought and elevated atmospheric CO_2_ to mature oak forests have differing impacts on the litter decomposition process. Specifically, our findings show that decomposition environment was the stronger force shaping mass loss at O_3_HP, which supports our first prediction. The stronger influence of litter origin on mass loss in the elevated CO_2_ experiment at BIFoR FACE provides support for our second prediction, highlighting the importance of carry over effects on leaves prior to decomposition.

### (i) Litter properties and their legacy effects

We found that litter incubated under trees subjected to long-term drought at O_3_HP had decomposed more slowly compared to litter incubated in ambient rainfall conditions. But critically, the C:N ratio, leaf litter functional groups (at the litterbag-level), and the composition of secondary metabolites differed, in part, according to the origin of the litter. Our finding that litter origin, capturing treatment-driven changes in initial litter continued months after the onset of decomposition suggests strong carry-over effects of global change factors acting *via* modifications on traits of living leaves. Changes in leaf traits, and thus the resulting litter traits, have been demonstrated in response to perturbations, such as nitrogen enrichment (Gill et al., 2022; Ping et al., 2025; Zhao et al., 2024), that impacts decomposition. Leaf and litter traits will influence the activity, biomass and community composition of soil food webs (Bourget et al., 2023; Fujii et al., 2018; Ni et al., 2025; Ridgeway et al., 2022) and alter products of litter decomposition that can become protected, like the formation of mineral associated organic matter (Jia & Feng, 2026). We also found that the carry-over effect of litter origin was reflected by the colonization and establishment of microbial communities (drought experiment) and animal abundances (elevated CO_2_ experiment) in litter. Specifically, microbial biomass (measured as total PLFA concentration) was higher in the experimental litter under drought, while fewer animals were present in experimental litter in response to elevated CO_2_. Overall, leaves grown under global change factors prior to litterfall continue to impact litter properties and decomposers even when >50% of litter has been decomposed. These results highlight the longevity of impacts of global changes on litter decomposition, and thus C cycling within forests.

Further to these carry-over effects, we found an effect of litter origin treatment in the elevated CO_2_ experiment on litter mass loss even though there were no significant differences in several initial litter properties known to regulate decomposition (Prescott, 2010), such as C:N and metabolites. This finding suggests that the treatment may have changed other important litter traits that impacted mass loss, despite small differences with initial litter C:N. Such effects could include increasing leaf and cuticle thickness, stemming from cell expansion under elevated CO_2_ (Pritchard et al., 1999) as greater thickness can slow litter decomposition (Castillo-Figueroa & Posada, 2025; Zukswert & Prescott, 2017), or changes to nutrient concentrations; previous work has shown that worsening nutritional status in leaves, such as declining phosphorus, potassium and micronutrient concentrations, can stem from elevated CO_2_ concentrations (Penuelas et al., 2020). Reduced nutrient concentrations in foliar tissue, besides carbon and nitrogen, may have decreased litter decomposition rates and impacted decomposer communities in our study (Kaspari et al., 2008; Kerekes et al., 2013). Future studies should expand on the leaf and litter traits that were measured to better understand the ways that elevated CO_2_ can impact leaf decomposition. A potential interaction between leaf physical and chemical traits influencing mass loss under elevated CO_2_ highlights the complexity by which global change factors will influence carbon and nutrient cycling in forests.

Analysis of litter properties using FTIR provided a more detailed perspective of how decomposition shapes litter functional groups, which enables better understanding of how decomposition may influence the formation of specific C pools with different rates of turnover, which underpins recent concepts of soil C cycling (e.g., Lavallee et al., 2020). In the drought experiment, FTIR analysis revealed strong differences between decomposition environment (subplot and litterbag-level) and litter origin (litterbag-level only), while few differences were detected between litter origins in the elevated CO_2_ experiment, mirroring the results from litter nutrient stoichiometry (C and N). Strong correlations between FTIR and other measurements have been found (Liu et al., 2016), but FTIR can also provide additional information on degradation of leaf litter material and whether litter properties converge (Moore et al., 2011) or diverge (Wickings et al., 2012) during mass loss. How litter functional groups change during decomposition will likely depend on interactions between time decomposed, initial litter properties, decomposer community, and environmental conditions. Overall, we found that FTIR functional groups in litter and mass loss responded similarly to treatments within each experiment, thus providing an additional methodology to consider when detailing how litter properties are reshaped by global change factors.

### (ii) Environmental impacts on decomposition

Amplified drought conditions at O_3_HP strongly shaped litter properties, decomposers, and mass loss. Drought has been shown repeatedly to slow decomposition rates across several ecosystems and climates (Liu et al., 2024) in addition to decreasing animal abundances on remaining litter (Martin et al., 2024) and shifting microbial communities to favour fungi (Canarini et al., 2024; de Vries et al., 2018; Yuste et al., 2011). In our study, we found drought slowed decomposition (Quer et al., 2022; Santonja et al., 2015), increased F:B ratios (Biryol et al., 2024), and modestly decreased mesofauna abundances (Santonja et al., 2017), aligning with prior research at the site and adjacent experiments. Although not statistically significant, there was a trend for decreasing oribatid mite abundance under drought, also consistent with previous studies (Aupic-Samain et al., 2021). Together, our results add further evidence to the pervasive effect of drought on litter mass loss. Importantly, they also highlight that the effect of droughted conditions on litter mass loss rates exceeds that of long-term carry over effects.

By contrast, we found that an enriched CO_2_ environment had no impact on mass loss or decomposer communities, and comparing with another elevated CO_2_ experiment in mature forest, EucFACE (e.g., Jiang et al., 2020), highlights apparent differences in environmental context. At EucFACE, elevated CO_2_ concentrations led to modest increases in decomposition rates, particularly at later decay stages (Nielsen et al., 2024) as impacts on fungal communities on litter were apparent. Other studies have also reported faster decomposition rates in enriched CO_2_ environments (Cotrufo et al., 2005; Hall et al., 2006), suggesting higher CO_2_ levels can increase mass loss rates in some scenarios, which may reflect changes to fungal community composition and activity (Carney et al., 2007; Ochoa-Hueso et al., 2017) and priming and faster turnover of soil carbon (Drake et al., 2011; Jiang et al., 2020). The contrasting results between our study and others highlights the context-dependency in which elevated CO_2_ environments impacts litter decomposition, and support a recent meta-analysis showing that elevated CO_2_ has a weak, inconsistent effect on mass loss (Liu et al., 2024). In summary, the decomposition environment under elevated CO_2_ was not a strong factor in shaping mass loss, litter properties and decomposers during decomposition.

### (iii) Microbial and faunal contributions to decomposition processes

Microbial communities (measured with PLFAs) from each experimental site were remarkedly different from one another, irrespective of the experimental manipulation, and likely had a strong influence on the relative rates of decomposition between sites. For example, F:B ratio was considerably higher at O_3_HP relative to BIFoR FACE, and this difference is likely explained by the overall higher C:N ratios of the initial litter at O_3_HP. Fungal abundance is typically greater on recalcitrant litter because fungi possess enzymes that breakdown larger carbon-rich molecules and polymers, such as lignin, relative to bacteria (Leifheit et al., 2024; Romaní et al., 2006; Wang & Kuzyakov, 2024), and litter C:N is often strongly related to microbial communities, including F:B ratio (Kohl et al., 2021).

Our findings showed that the decomposition environment under drought was the primary control of microbial communities, with the litter origin having lesser effects. Microbial communities are often tied to decomposition and the impacts of drought on them may be a key control of decomposition. For example, Allison et al. (2013) found that litter inoculated with microbes from droughted plots decreased mass loss rates under ambient moisture conditions demonstrating that both the drought environment and microbial origin influenced decomposition. Long-term drought manipulations may favour drought-tolerated taxa leading to changing functional responses, such as decreasing enzyme production and increasing energy expenditure in survival which may impact mass loss rates (Allison & Goulden, 2017; Canarini et al., 2024; Malik et al., 2020). Further manipulation experiments are needed to determine whether long-term (10+ years) drought at O_3_HP has selected for drought tolerant microbial communities.

The lack of significant changes to microbial communities at BIFoR FACE is surprising given the numerous microbially-driven, below-ground impacts reported at this site. These effects include increases in root exudation rates (Rumeau et al., 2025), root traits, such as branching frequency (Reay et al., 2025) and the abundance and turnover of mycorrhizal hyphae. There are two hypotheses that might explain these findings. First, the microorganisms associated with freshly fallen litter may be influenced more by the litter itself than the surrounding soil (decomposition) environment. For example, there is evidence that bulk soil and litter can harbour distinct microbial communities, with the latter driven by species identity and leaf traits (Prescott & Grayston, 2013). Second, many ectomycorrhizal fungi proliferate in the fermentation and humic horizons i.e. acting on partially degraded litter (Dickie et al., 2002; Rosling et al., 2003), and may be more closely associated with roots (Cahanovitc et al., 2022) and not freshly fallen litter. In general, it is unclear to what extent broad changes in ectomycorrhizal hyphal production previously measured in response to elevated CO_2_ at BIFoR FACE (Reay et al., 2025) impacted our results. For example, some specific ectomycorrhizal fungal taxa can accelerate decomposition of relatively fresh litter (Lindahl et al., 2021), while the activity and abundance of many ectomycorrhizal fungi often suppress litter decomposition through the well-known ‘Gadgil’ effect (Choreño Parra & Treseder, 2024) and may have impacted mass loss rates in our study.

We found that experimental litter from the elevated CO_2_ experiment and litter incubated under drought conditions were populated with fewer mesostigmatid and prostigmatid mites. Most mesostigmatid mites, and some prostigmatid mite species, are predators, feeding on arthropods and nematodes (Lindquist et al., 2009; Walter et al., 2009). Lower abundance of these mites suggests that changes to prey composition (but likely not group abundances in our experiments) hindered predators in our litterbags. Changing mesofauna and macrofauna predator composition can indirectly influence litter decomposition and soil nutrient fluxes (Koltz et al., 2018; Staddon et al., 2010). For example, macrofauna predators can reduce competition amongst lower trophic levels, resulting in higher litter decomposition, particularly in dry forests (Melguizo-Ruiz et al., 2020) as the effects of top-down control by predators, and its indirect effects on litter decomposition can depend on moisture availability and temperature (Lang et al., 2014; Lawrence & Wise, 2004). Soil animals consistently increase decomposition, even under drought (Luan et al., 2024), thus changing trophic dynamics under global change factors may directly or indirectly impact mass loss rates.

Our experimental design enabled us to test for interactive effects between litter origin and decomposition environment on decomposer communities. This analysis is important because litter properties and decomposers interact during decomposition (Prieto et al., 2019) and microbes and animals often show a consistent ability to accelerate decomposition of ‘home’ litter relative to ‘away’ litter (i.e. the home field advantage) (Ayres et al., 2009; Veen et al., 2015). This variation is driven partly by litter quality (e.g., C:N ratio), where recalcitrant litter is typically subject to stronger home field advantage than more labile litter (Chomel et al., 2015; Wallenstein et al., 2013). Potentially, changing decomposer communities alongside litter properties due to drought or elevated CO_2_ may lead to synergistic (or antagonistic) responses to mass loss. Nevertheless, there was no significant interaction between the litter origin and decomposition environment treatments with litter mass loss under either treatment, suggesting that changes to the decomposer community or initial litter properties under droughted and elevated CO_2_ conditions was not strong enough to produce an interactive response.

### (iv) Resolving mechanisms across experiments: caveats and future studies

Our findings are derived from two large-scale, long-term manipulations of climate change factors in ancient oak woodland. Deploying and analyzing litterbags using a common experimental design enabled us to rigorously test how elevated CO_2_ and drought affect decomposition through two pathways, the decomposition environment and initial litter properties. Thus, over-coming limitations of equivalent analyses undertaken in meta-analyses where data are invariably derived using markedly different approaches. The caveat to our analysis is that we were constrained to just two locations (unlike most meta-analyses), with one location for each climate change factor, and have limited power to test how the background environmental conditions, litter source characteristics and soil properties interact with treatments. Interactions between climate with litter properties, biota, and decomposition are well established (Canessa et al., 2021; García-Palacios et al., 2013; Glassman et al., 2018) and these interactions will impact how global change factors impact decomposition. Thus, our findings are limited by the fact we cannot fully disentangle the role of treatments from that of local environmental conditions. However, our results reveal several outcomes that can be expanded in future research to examine the impact of global change factors on litter decomposition. First, we highlight the strength of employing a common experimental design across contrasting climates to understand how local climate and soil properties influence litter decomposition under global change factors (e.g., Nielsen et al., 2024). Second, our findings demonstrate a need to consider a greater range of physical and chemical leaf trait measurements to provide greater explanatory power to explain decomposition (e.g., Pichon et al., 2020). Third, parameterizing microbial driven decomposition from that of animals (e.g., Luan et al., 2024) in addition to considering contrasting litter types (e.g., deciduous leaves vs. coniferous needles (e.g., Yang et al., 2022) to identify how decomposers influence litter types and its interactions with global change factors. Finally, incorporating other soil properties, pools, and functions, such as mineral associated organic carbon (e.g., Sokol et al., 2026), to investigate the broader impacts of global change factors on litter decomposition. Together, these approaches will advance our collective understanding of how global change factors shape litter decomposition.

## Conclusions

Our findings reveal how separate *in situ* manipulation of elevated CO_2_ or drought impact litter decomposition through distinct pathways involving changes in either litter properties prior to decomposition or the decomposition environment itself. Consistent with our predictions, drought exerted the strongest control on decomposition, primarily through environmental constraints such as reduced soil moisture and altered microclimatic conditions, while changes to mass loss rates under in enriched CO_2_ environments was found in the litter origin treatment. We also found clear evidence that the impact of drought and elevated CO_2_ leads to strong carry-over effects on litter properties and biota, confirming the importance of pre-litterfall changes in initial litter alongside *in situ* environmental modification.

The findings presented here have broad implications for litter decomposition under global changes factors. In the future, increasing annual atmospheric CO_2_ concentrations may alter litter decomposition through alterations to litter properties, which may include C:N ratio, underlying litter chemical functional groups but also other leaf physiological properties, as changes to the decomposition environment may weakly alter mass loss rates. In conjunction, more frequent and intense droughts under climate change will strongly shape litter properties prior to litterfall and during decomposition, alongside the associated decomposer communities. Overall, our findings, using long-term, realistic *in situ* manipulations, provide a more complete understanding of how global change factors shape C cycling in forests. Our experiment highlights that litter decomposition will be reshaped by global change factors through multiple pathways as further research should uncover the broader impacts of drought and elevated CO_2_ on forest and soil systems.

## Supporting information

Supplmentary information for the main texts containing additional tables, figures, and information

## Acknowledgments

This study was funded by a UKRI Natural Environment Research Council (NERC) grant NE/S002189/1. ARMK gratefully acknowledges additional support from NERC through grant NE/S015833/1 (QUINTUS). TC acknowledges support by Research Ireland (20/FFP-P/8584). The BIFoR-FACE facility acknowledges generous underpinning support from the JABBS Foundation, Norbury Park Estate, The John Horseman Trust, Ecological Continuity Trust, and the University of Birmingham. We thank Dr Kris Hart and the operations team at BIFoR FACE, Dr. Ully Kritzler, Deborah Ashworth, Dr. Thomas Bishop, and Dr. Benjamin Bell for technical support. We thank Dr. Stéphane Greff (Aix-Marseille Université) for analyzing leaf litter and Jean-Phillipe Orts (Aix-Marseille Université) for collecting litter from the O_3_HP experimental site.

## Data availability statement

Data will be submitted to Figshare upon acceptance of the manuscript.

## Author Contributions

DJ and MC conceptualized the study, and DJ, MC, VB, RDB, ARMK obtained funding. MLM, MC, ZV and KJF collected data. ARMK and VB help designed the elevated CO_2_ and drought sites, respectively, and coordinated work there. MLM and MC curated data, and MLM performed the data analysis with support from TC. MLM wrote the first version of the manuscript and MC, KJF, TC, ARMK, VB, RDB, and DJ provided feedback and contributed to finalizing the manuscript.

