## Supplementary material for "Global change factors reshape the links between litter properties, decomposers, and decomposition in mature oak forests": Supplmentary information for the main texts containing additional tables, figures, and information

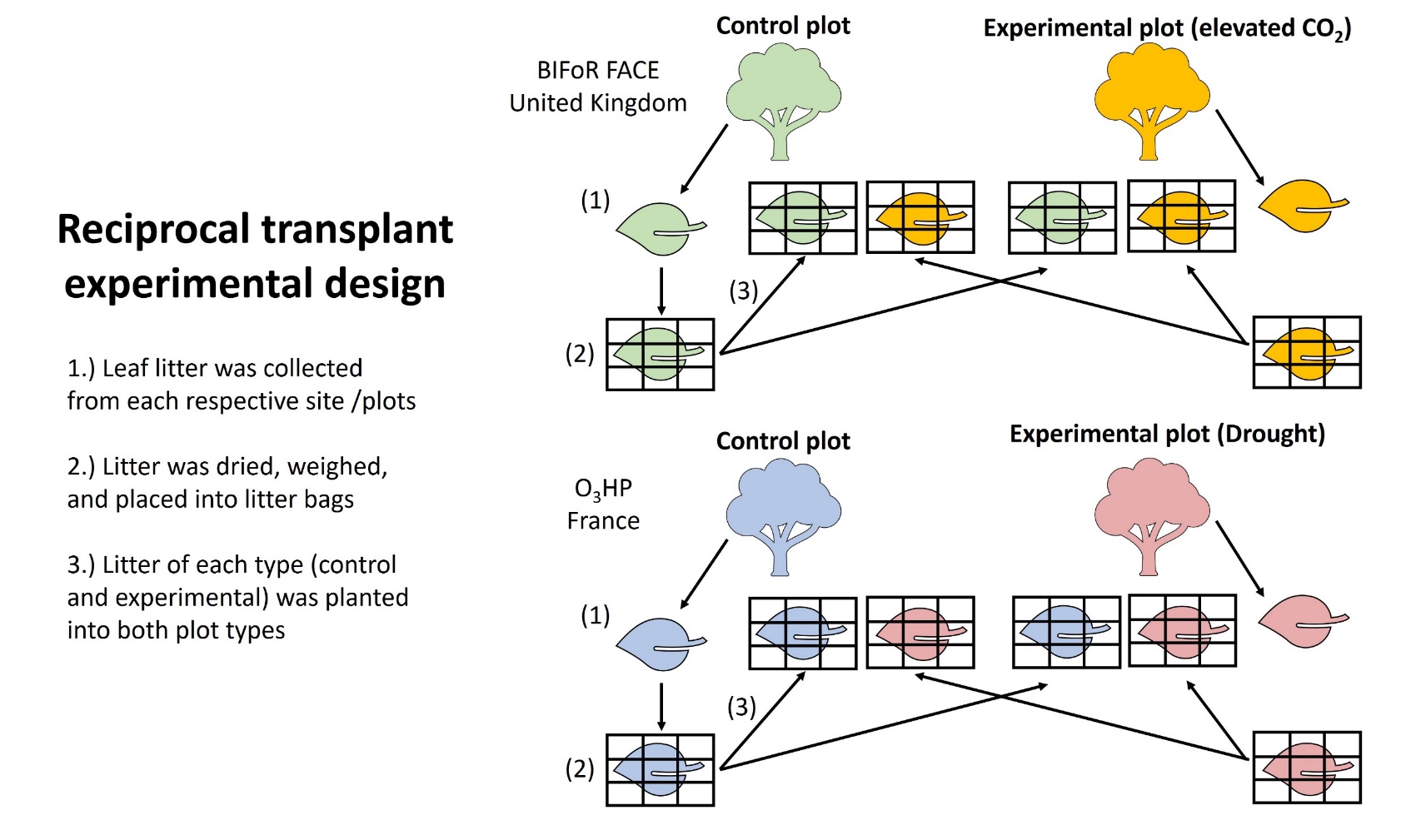

Fig. S1 Overview of our litterbag study that used a reciprocal transplant experimental design. Two separate independent experiments were conducted: in one litter was collected from control and droughted plots at the Oak Observatory at the Observatoire de Haute-Provence (O_3_HP), and in another from control and elevated CO_2_ [eCO_2_] plots at the Birmingham Institute of Forest Research Free-Air CO_2_ Enrichment (BIFoR FACE). Litter was placed into 1800 micrometre-mesh litterbags and transplanted back into both control and experimental areas to decompose and collected over four time points (i.e., harvests).

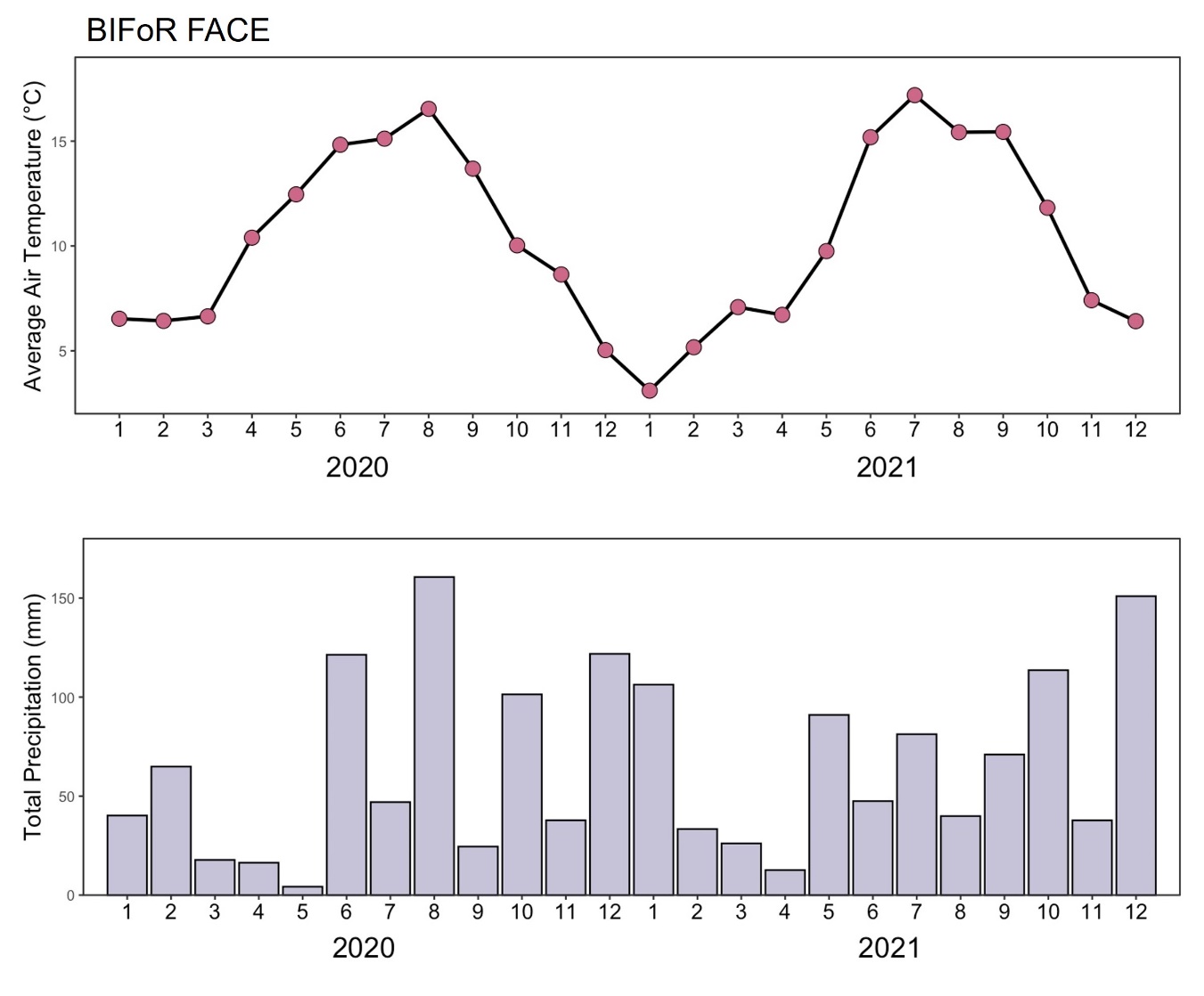

Fig. S2 Monthly average temperature and total precipitation at the BIFoR FACE site during the study. The x-axis shows year and month (labelled as 1-12 which correspondence to months January to December, in that order). Litterbags were placed into respective plots January 2020 and harvested across four time points: September 2020 (harvest one (H1)), January 2021 (harvest two (H2)), June 2021 (harvest three (H3)) and December 2021 (harvest four (H4)).

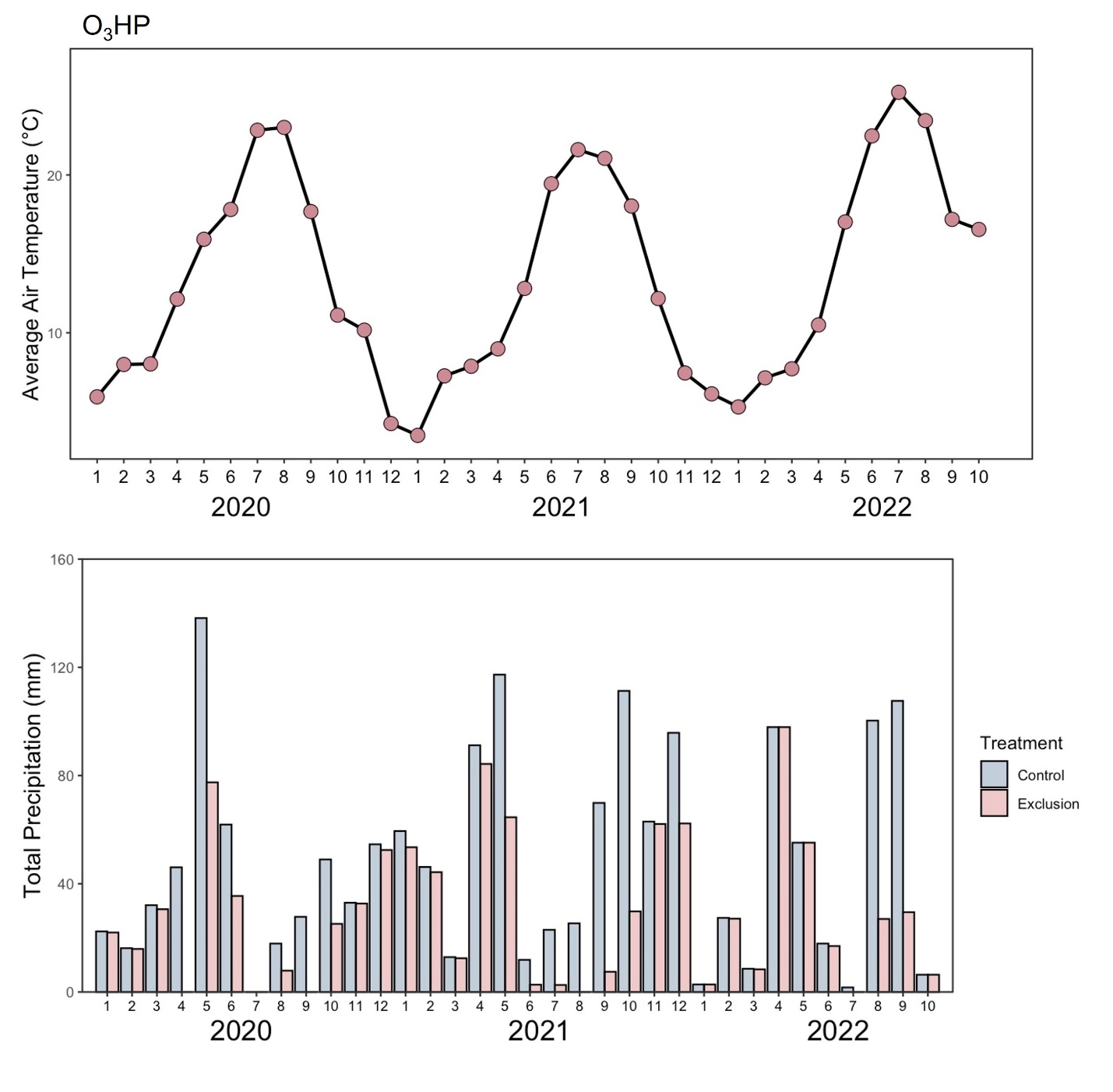

Fig. S3 Monthly average temperature and total precipitation at the O_3_HP site during the study. The x-axis shows year and month (labelled as 1-12 which correspondence to months January to December, in that order). Litterbags were placed into respective plots January 2020 and harvested across four time points: October 2020 (harvest 1 (H1)), January 2021 (harvest 2 (H2)), September 2021 (harvest 3 (H3)) and October 2022 (harvest 4 (H4)).

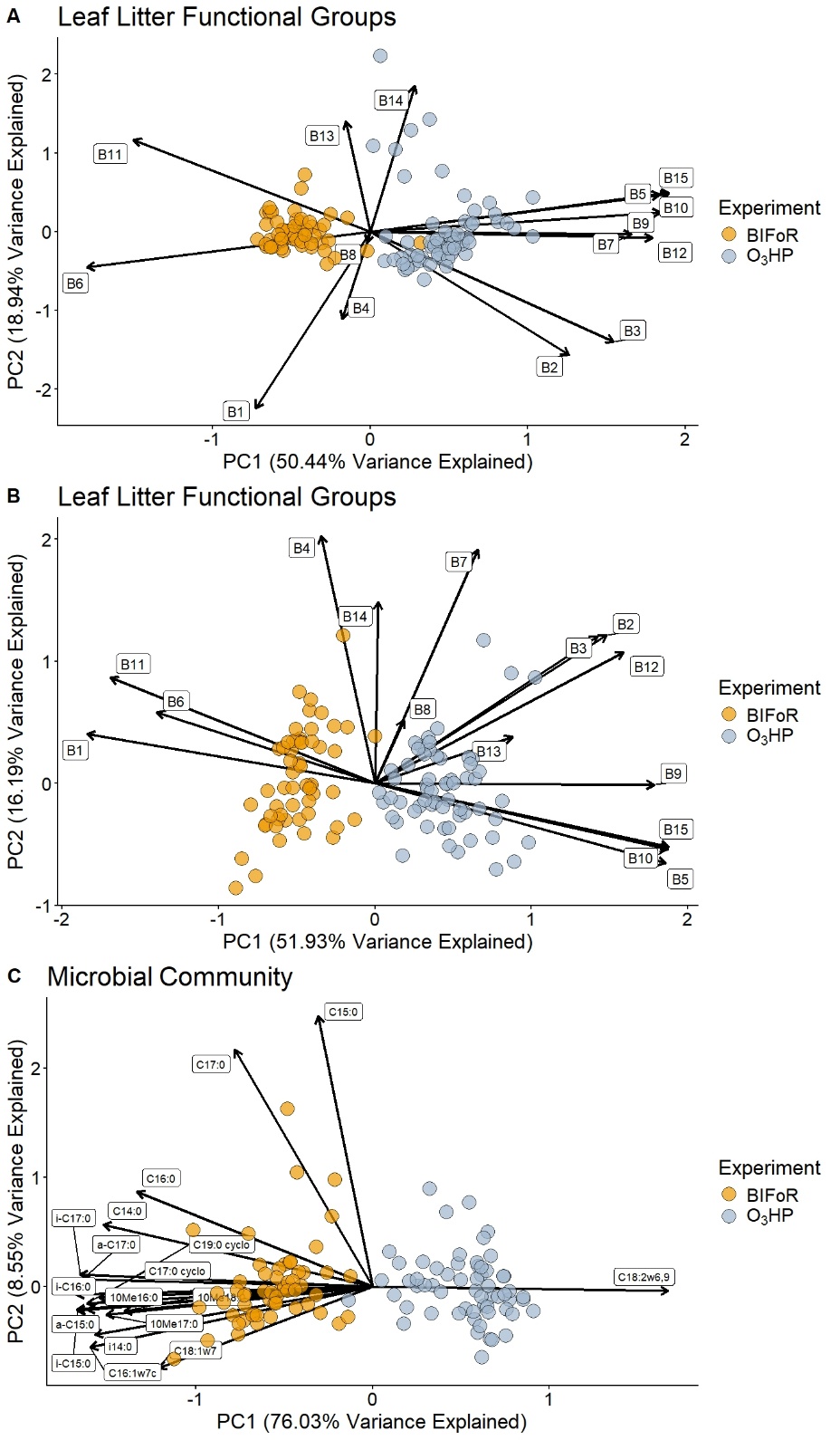

Fig. S4 Principal component analysis (PCA) ordinations of leaf litter functional groups measured by Fourier transform infrared (FTIR) spectroscopy at the third (A) and fourth (B) harvest. C) Principal component analysis (PCA) ordination of microbial community structure measured by phospholipid fatty acid analysis (PLFA) at the third harvest. Points are coloured by the respective experiments, BIFoR FACE (elevated CO_2_) and O_3_HP (drought).

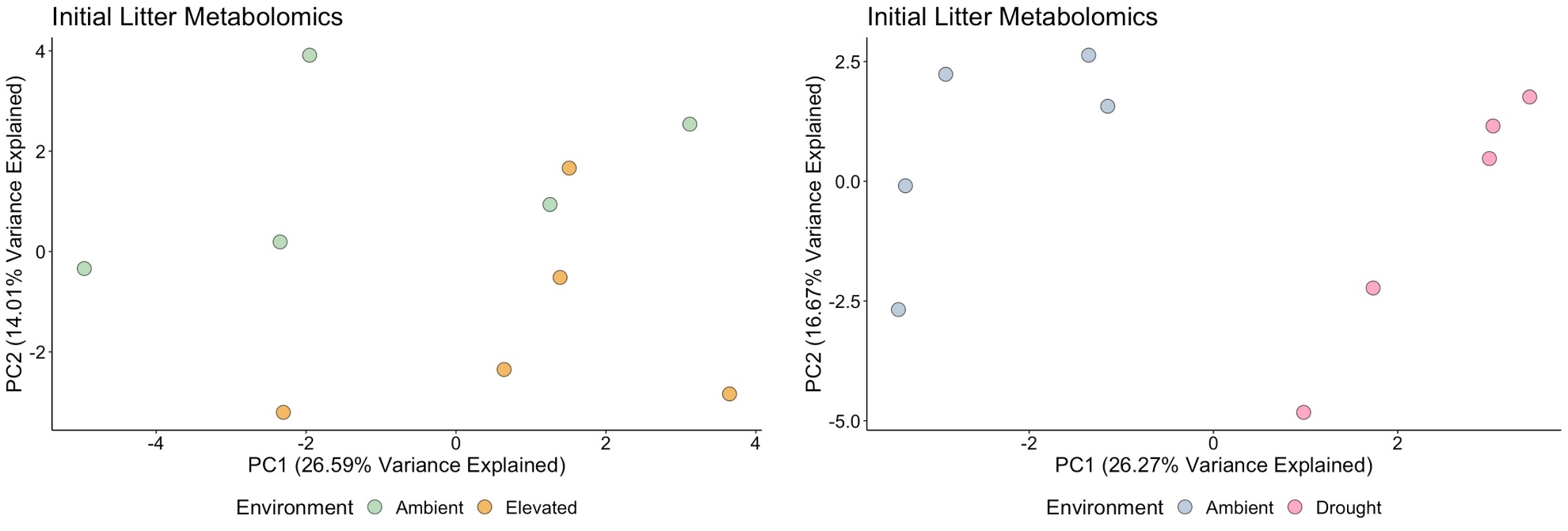

Fig. S5 Principal component analysis (PCA) of the initial litter metabolomic profile (where values were log10(x+1), z-transformed) for litter from the elevated CO_2_ experiment (left) from BIFoR FACE and the drought experiment (right) from O_3_HP. Using PERMANOVAs (on log10+1, z-transformed data), we found strong differences between the metabolomic profiles of the initial control and droughted litter (R_2_ = 0.25, F_1,8_ = 2.62, p = 0.009) but not between ambient and elevated CO_2_ litter (R_2_ = 0.14 F_1,8_ = 1.34, p = 0.116) although differences in metabolomics can be seen visually in the PCA for the elevated CO_2_ experiment.

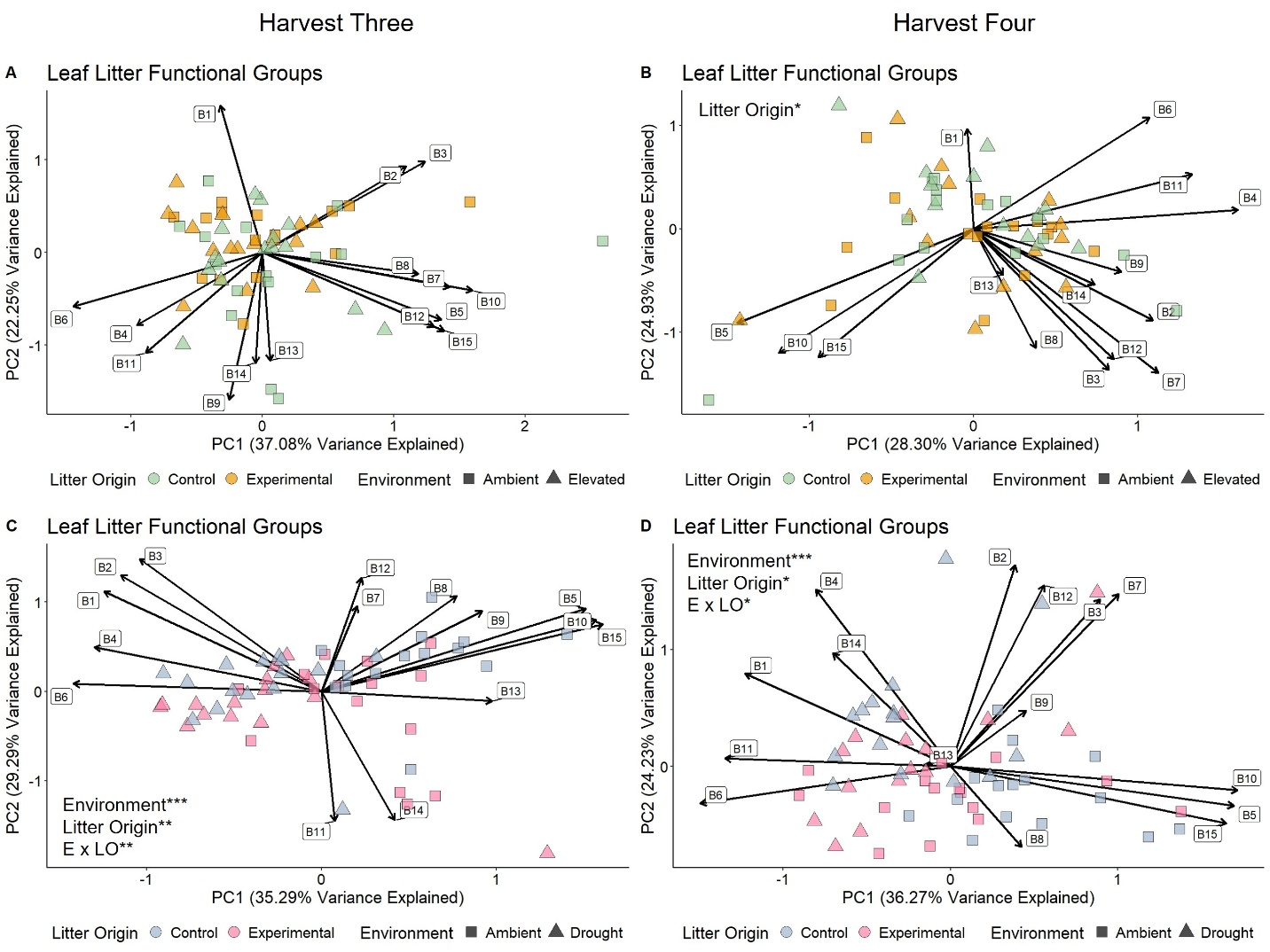

Figure S6. Principal component analysis (PCA) ordinations of leaf litter functional groups measured by Fourier transform infrared (FTIR) spectroscopy for the elevated CO_2_ experiment (top) and drought experiment (bottom) at harvest three (A and C, left) and four (B and D, right). Points represent individual litterbags. Corresponding functional groups for band numbers (B1, B2, B3, etc.) can be found in Supplementary Information Table S1. * = p < 0.05, ** = p < 0.01, and *** = p < 0.001. We have only provided p-values of factors that were statistically significant. E = Environment and LO = litter origin.

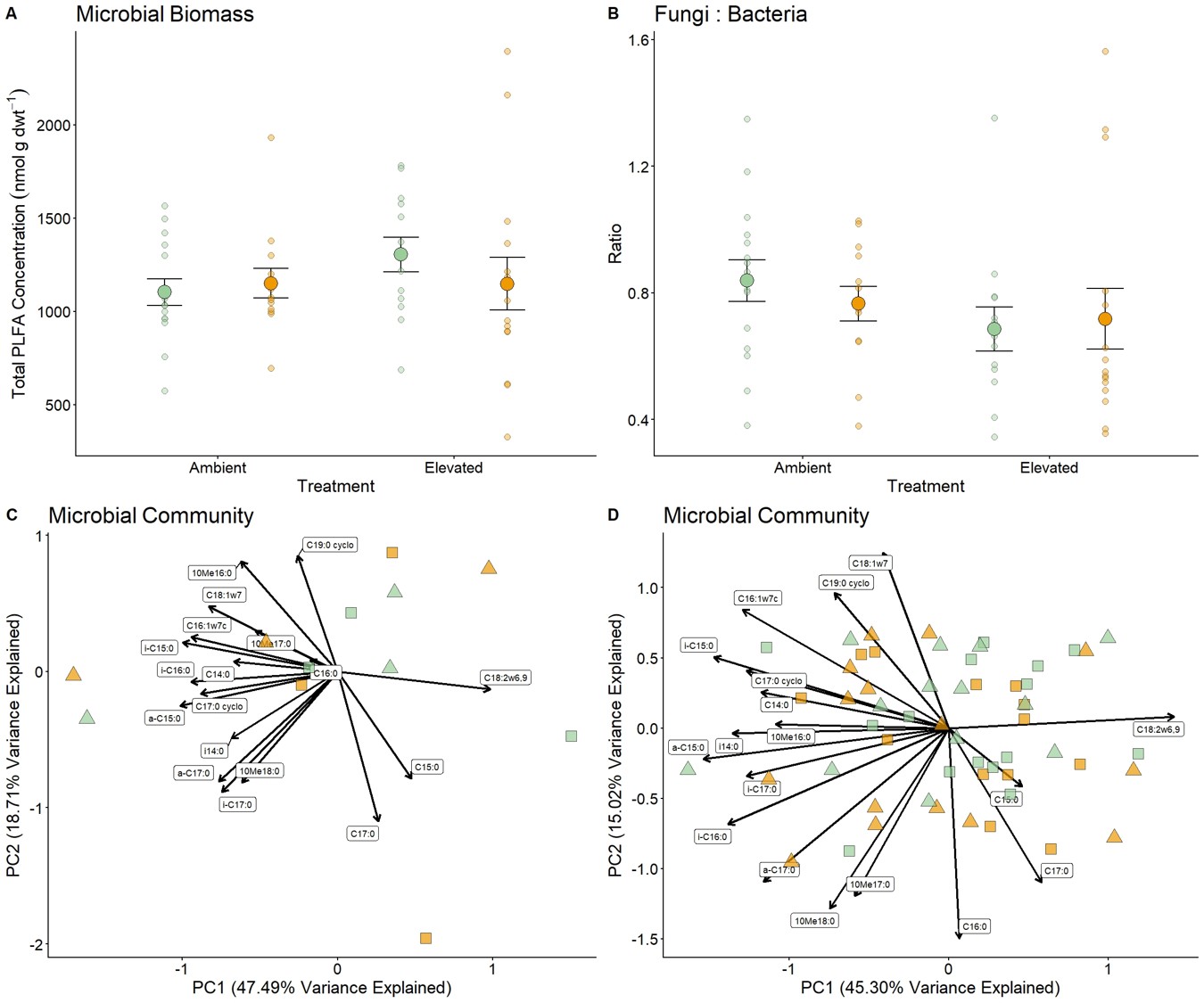

Fig. S7 Phospholipid fatty acid analysis (PLFA) composition of the microbial community in the elevated CO_2_ experiment at H3. For A) (total PLFA) and B) (fungi : bacteria) larger points represent mean (± SE) while smaller points are the raw data values for individual litter bags. Principal component analysis (PCA) ordination of microbial community structure at subplot-level (C) and litterbag-level (D). Neither litter origin nor decomposition environment significantly affected PLFA concentration, fungi : bacteria., or composition.

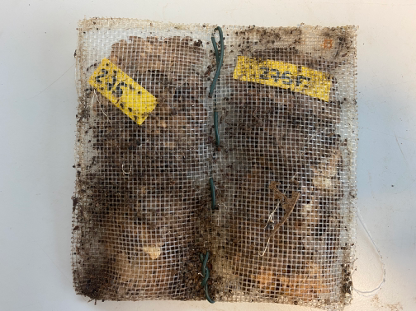

Image S1. Example of harvested litter bag (litter bag 275) from the O_3_HP site (drought experiment). Note the two compartments, one labelled 275 and the other 275F used for chemical and faunal analysis, respectively.

Table S1 Band numbers and regions of assigned litter functional groups using Fourier transform infrared (FTIR) spectroscopy. Assignment to functional group was based on Liu et al., (2016).

| Band number | Wave number region (cm^-1^) | Functional group |
| --- | --- | --- |
| B1 | 3689-3033 | hydroxyl groups |
| B2 | 2995-2881 | aliphatic C-H-stretch |
| B3 | 2865-2834 | C-H-stretch |
| B4 | 1783-1708 | amide I, C==0-stretch |
| B5 | 1696-1550 | C==C, carboxylates |
| B6 | 1536-1448 | N-H, C==C, amide II |
| B7 | 1487-1441 | aliphatic groups |
| B8 | 1441-1402 | amide, carboxylates |
| B9 | 1397-1349 | acetyl groups |
| B10 | 1349-1301 | OH, sulfones, esters |
| B11 | 1301-1196 | OH,==C-O-C, phenols |
| B12 | 1184-1143 | aliphatic OH, ethers |
| B13 | 1143-940 | polysaccharides, C-OH |
| B14 | 907-885 | carbohydrates |
| B15 | 798-770 | aromatic systems |

Table S2 Summary table showing the mean (± SE) of %C, %N and C:N ratio of initial litter collected from five samples. Litter was collected October 2019 from BIFoR (elevated CO_2_ experiment) and O_3_HP (drought experiment) where it was then air dried, ball-milled, prior to being analyzed on a Elementar Vario EL element analyser. Bolded averages equate to significant differences between litter types and parameters.

| Site / Treatment | Litter Quality Indices | | |
| --- | --- | --- | --- |
| BIFoR FACE | %C | %N | C:N |
| Ambient CO_2_ | 45.9 ± 0.91 | 1.2 ± 0.02 | 39.9 ± 1.14 |
| Elevated CO_2_ | 46.8 ± 0.28 | 1.2 ± 0.06 | 39.8 ± 1.98 |
| O_3_HP | %C | %N | C:N |
| Control | **45.0 ± 0.19** | **0.6 ± 0.02** | **73.0 ± 1.8** |
| Drought | **45.8 ± 0.21** | **0.8 ± 0.04** | **60.2 ± 2.9** |

Table S3 Summary table of mean (± SE) of the exponential decomposition rate coefficient (*k*) for BIFoR FACE (elevated CO_2_ experiment, top) and O_3_HP (drought experiment, bottom) across all four harvests.

**BIFoR FACE**

| Harvest | Environment | Litter Origin | *k* |
| --- | --- | --- | --- |
| 1 | Ambient | Control | 1.00 ± 0.05 |
|  |  | Experimental | 0.90 ± 0.04 |
|  | Elevated CO_2_ | Control | 1.03 ± 0.07 |
|  |  | Experimental | 0.92 ± 0.10 |
| 2 | Ambient | Control | 1.02 ± 0.07 |
|  |  | Experimental | 0.94 ± 0.07 |
|  | Elevated CO_2_ | Control | 0.96 ± 0.04 |
|  |  | Experimental | 0.95 ± 0.07 |
| 3 | Ambient | Control | 0.74 ± 0.05 |
|  |  | Experimental | 0.69 ± 0.04 |
|  | Elevated CO_2_ | Control | 0.88 ± 0.06 |
|  |  | Experimental | 0.77 ± 0.06 |
| 4 | Ambient | Control | 0.89 ± 0.04 |
|  |  | Experimental | 0.90 ± 0.07 |
|  | Elevated CO_2_ | Control | 0.90 ± 0.05 |
|  |  | Experimental | 0.92 ± 0.09 |

**O_3_HP**

| Harvest | Environment | Litter Origin | *k* |
| --- | --- | --- | --- |
| 1 | Ambient | Control | 0.42 ± 0.03 |
|  |  | Experimental | 0.46 ± 0.02 |
|  | Drought | Control | 0.29 ± 0.01 |
|  |  | Experimental | 0.32 ± 0.01 |
| 2 | Ambient | Control | 0.59 ± 0.02 |
|  |  | Experimental | 0.60 ± 0.02 |
|  | Drought | Control | 0.49 ± 0.02 |
|  |  | Experimental | 0.51 ± 0.03 |
| 3 | Ambient | Control | 0.43 ± 0.02 |
|  |  | Experimental | 0.45 ± 0.01 |
|  | Drought | Control | 0.31 ± 0.01 |
|  |  | Experimental | 0.32 ± 0.01 |
| 4 | Ambient | Control | 0.39 ± 0.01 |
|  |  | Experimental | 0.39 ± 0.02 |
|  | Drought | Control | 0.28 ± 0.01 |
|  |  | Experimental | 0.30 ± 0.03 |

Table S4 Summary table showing mean (± SEM) of decomposition environment and litter origin for measured univariate parameters for harvest three in the elevated CO_2_ experiment (BIFoR FACE) along with the results to linear mixed models: * = p < 0.05, ** = p < 0.01, and *** = p < 0.001. R^2^m = marginal R^2^. R^2^c = conditional R^2^. If no marginal R^2^ is provided, it is due to the random factor being removed from the model (stemming from singularity with the random factor). In this case, the adjusted R^2^ is then provided as R^2^c.

| Environment (E) | Ambient | | Elevated CO_2_ | |  |  |  |  |  |
| --- | --- | --- | --- | --- | --- | --- | --- | --- | --- |
| Litter Origin (LO) | Control | Experimental | Control | Experimental | E | LO | E x LO | R^2^m | R^2^c |
| Moisture Content (%) | 49.0 ± 3.59 | 51.2 ± 4.07 | 60.0 ± 3.34 | 52.8 ± 3.96 |  |  |  | 0.075 | 0.160 |
| %C | 45.9 ± 0.48 | 46.6 ± 0.33 | 46.2 ± 0.28 | 46.6 ± 0.39 |  |  |  | 0.040 | 0.120 |
| %N | 2.2 ± 0.02 | 2.2 ± 0.04 | 2.2 ± 0.02 | 2.1 ± 0.02 |  |  |  |  | 0.009 |
| C:N | 21.0 ± 0.27 | 21.8 ± 0.43 | 21.0 ± 0.24 | 21.8 ± 0.32 |  | * |  |  | 0.050 |
| PLFA concentration  (nmol g dwt^-1^) | 1103 ± 71.2 | 1151 ± 79.6 | 1305 ± 92.4 | 1148 ± 141.8 |  |  |  |  | -0.016 |
| Fungi : Bacteria | 0.83 ± 0.07 | 0.77 ± 0.05 | 0.69 ± 0.07 | 0.72 ± 0.10 |  |  |  | 0.044 | 0.178 |
| Collembola Abundance (individuals g dwt^-1^) | 69.5 ± 11.22 | 76.8 ± 11.9 | 82.7 ± 9.4 | 82.3 ± 10.68 |  |  |  | 0.015 | 0.191 |
| Oribatida Abundance (individuals g dwt^-1^) | 56.2 ± 6.67 | 50.8 ± 8.22 | 57.4 ± 7.62 | 63 ± 9.61 |  |  |  | 0.018 | 0.164 |
| Mesostigmata Abundance (individuals g dwt^-1^) | 23.0 ± 2.94 | 21.5 ± 3.16 | 26.0 ± 3.32 | 26.5 ± 2.91 |  |  |  | 0.029 | 0.115 |
| Astigmata Abundance (individuals g dwt^-1^) | 1.1 ± 0.33 | 0.8 ± 0.27 | 1.3 ± 0.39 | 1.9 ± 0.47 |  |  |  | 0.068 | 0.143 |
| Prostigmata Abundance (individuals g dwt^-1^) | 14.7 ± 2.37 | 19.0 ± 2.27 | 16.4 ± 2.16 | 13.2 ± 2.21 |  |  |  | 0.057 | 0.087 |
| ‘Other’ Fauna Abundance (individuals g dwt^-1^) | 5.3 ± 0.78 | 5.8 ± 0.94 | 17.6 ± 5.61 | 10.9 ± 2.23 |  |  |  | 0.120 | 0.147 |
| Total Animal Abundance (individuals g dwt^-1^) | 169.9 ± 18.14 | 174.6 ± 23.07 | 201.3 ± 20.35 | 197.7 ± 22.26 |  |  |  | 0.027 | 0.220 |

Table S5 Summary table showing mean (± SEM) of decomposition environment and litter origin for measured univariate parameters for harvest four in the elevated CO_2_ experiment (BIFoR FACE) along with the results to linear mixed models: * = p < 0.05, ** = p < 0.01, and *** = p < 0.001. R^2^m = marginal R^2^. R^2^c = conditional R^2^. If no marginal R^2^ is provided, it is due to the random factor being removed from the model (stemming from singularity with the random factor). In this case, the adjusted R^2^ is then provided as R^2^c.

| Environment (E) | Ambient | | Elevated CO_2_ | |  |  |  |  |  |
| --- | --- | --- | --- | --- | --- | --- | --- | --- | --- |
| Litter Origin (LO) | Control | Experimental | Control | Experimental | E | LO | E x LO | R^2^m | R^2^c |
| Moisture Content (%) | 77.3 ± 0.94 | 78.1 ± 1.37 | 77.2 ± 0.95 | 77.5 ± 0.91 |  |  |  | 0.009 | 0.043 |
| %C | 46.8 ± 0.49 | 46.7 ± 0.52 | 46.9 ± 0.37 | 46.5 ± 0.47 |  |  |  | 0.006 | 0.021 |
| %N | 2.3 ± 0.04 | 2.3 ± 0.03 | 2.3 ± 0.05 | 2.3 ± 0.05 |  |  |  | 0.003 | 0.241 |
| C:N | 20.6 ± 0.43 | 20.5 ± 0.47 | 20.8 ± 0.47 | 20.5 ± 0.5 |  |  |  | 0.006 | 0.120 |
| Collembola Abundance (individuals g dwt^-1^) | 26.0 ± 2.08 | 24.9 ± 4.13 | 30.3 ± 5.74 | 25.3 ± 2.85 |  |  |  | 0.021 | 0.179 |
| Oribatida Abundance (individuals g dwt^-1^) | 70.7 ± 12.35 | 61.2 ± 8.39 | 62.3 ± 7.22 | 57.6 ± 4.95 |  |  |  |  | -0.035 |
| Mesostigmata Abundance (individuals g dwt^-1^) | 15.2 ± 1.18 | 11.6 ± 1.53 | 12.4 ± 1.48 | 9.9 ± 1.18 |  | * |  |  | 0.083 |
| Astigmata Abundance (individuals g dwt^-1^) | 3.2 ± 0.67 | 5.4 ± 2.13 | 5.6 ± 2.18 | 7.5 ± 2.99 |  |  |  | 0.012 | 0.347 |
| Prostigmata Abundance (individuals g dwt^-1^) | 13.7 ± 2.22 | 7.3 ± 1.26 | 8.9 ± 1.34 | 6.9 ± 0.83 |  | ** |  |  | 0.150 |
| ‘Other’ Fauna Abundance (individuals g dwt^-1^) | 5.4 ± 1.32 | 4.4 ± 0.73 | 4.4 ± 1.05 | 5.4 ± 1.28 |  |  |  |  | -0.041 |
| Total Animal Abundance (individuals g dwt^-1^) | 134.1 ± 13.30 | 114.7 ± 12.19 | 123.9 ± 13.6 | 112.5 ± 8.08 |  |  |  |  | -0.018 |

Table S6 Summary table showing mean (± SEM) of decomposition environment and litter origin for measured univariate parameters for harvest three in the drought experiment (O_3_HP) along with the results to linear mixed models: * = p < 0.05, ** = p < 0.01, and *** = p < 0.001. na = data was not analyzed due to low abundances. R^2^m = marginal R^2^. R^2^c = conditional R^2^. If no marginal R^2^ is provided, it is due to the random factor being removed from the model (stemming from singularity with the random factor). In this case, the adjusted R^2^ is then provided as R^2^c.

| Environment (E) | Ambient | | Drought | |  |  |  |  |  |
| --- | --- | --- | --- | --- | --- | --- | --- | --- | --- |
| Litter Origin (LO) | Control | Experimental | Control | Experimental | E | LO | E x LO | R^2^m | R^2^c |
| Moisture Content (%) | 56.7 ± 1.63 | 53.7 ± 2.72 | 34.9 ± 1.35 | 36.6 ± 1.19 | *** |  |  | 0.660 | 0.672 |
| %C | 44.7 ± 0.26 | 46.2 ± 0.12 | 47.3 ± 0.18 | 46.9 ± 0.18 | ** | ** | *** | 0.629 | 0.682 |
| %N | 1.2 ± 0.03 | 1.4 ± 0.03 | 1.1 ± 0.02 | 1.2 ± 0.02 | * | *** |  | 0.586 | 0.647 |
| C:N | 37.5 ± 1.07 | 32.8 ± 0.68 | 44.2 ± 0.99 | 39.4 ± 0.78 | ** | *** |  | 0.576 | 0.643 |
| PLFA Concentration  (nmol g dwt^-1^) | 1542.1 ± 53.1 | 1762.5 ± 60.8 | 1225 ± 55.9 | 1360.3 ± 75.4 | * | ** |  | 0.410 | 0.481 |
| Fungi : Bacteria | 3.1 ± 0.26 | 2.6 ± 0.14 | 3.9 ± 0.19 | 3.8 ± 0.18 | * |  |  | 0.332 | 0.358 |
| Collembola Abundance (individuals g dwt^-1^) | 25.4 ± 3.45 | 28.6 ± 4.18 | 17 ± 2.94 | 30.7 ± 7.43 |  |  |  | 0.072 | 0.164 |
| Oribatida Abundance (individuals g dwt^-1^) | 69.3 ± 7.68 | 78.8 ± 9.13 | 42.9 ± 6.34 | 49.1 ± 9.13 |  |  |  | 0.168 | 0.372 |
| Mesostigmata Abundance (individuals g dwt^-1^) | 10.7 ± 1.17 | 9.6 ± 0.70 | 4.5 ± 0.58 | 5.3 ± 1.11 | *** |  |  |  | 0.332 |
| Astigmata Abundance (individuals g dwt^-1^) | 1.2 ± 0.96 | 0.2 ± 0.07 | 0.2 ± 0.10 | 0.7 ± 0.31 | na | na | na | na | na |
| Prostigmata Abundance (individuals g dwt^-1^) | 31.9 ± 3.14 | 32.9 ± 2.85 | 31.2 ± 5.62 | 23.3 ± 2.55 |  |  |  |  | 0.020 |
| ‘Other’ Fauna Abundance (individuals g dwt^-1^) | 3.7 ± 0.48 | 4.1 ± 0.49 | 8 ± 0.81 | 5.4 ± 0.65 | * |  | * | 0.328 | 0.383 |
| Total Animal Abundance (individuals g dwt^-1^) | 142.1 ± 8.45 | 154.1 ± 13.2 | 103.8 ± 9.85 | 114.4 ± 15.31 |  |  |  | 0.154 | 0.330 |

Table S7 Summary table showing mean (± SEM) of decomposition environment and litter origin for measured univariate parameters for harvest four in the drought experiment (O_3_HP) along with the results to linear mixed models: * = p < 0.05, ** = p < 0.01, and *** = p < 0.001. na = data was not analyzed due to low abundances. R^2^m = marginal R^2^. R^2^c = conditional R^2^. If no marginal R^2^ is provided, it is due to the random factor being removed from the model (stemming from singularity with the random factor). In this case, the adjusted R^2^ is then provided as R^2^c.

| Environment (E) | Ambient | | Drought | |  |  |  |  |  |
| --- | --- | --- | --- | --- | --- | --- | --- | --- | --- |
| Litter Origin (LO) | Control | Experimental | Control | Experimental | E | LO | E x LO | R^2^m | R^2^c |
| Moisture Content (%) | 22.8 ± 3.6 | 21.6 ± 3.21 | 23.2 ± 2.79 | 27.3 ± 3.36 |  |  |  |  | -0.018 |
| %C | 45.1 ± 0.27 | 46.4 ± 0.27 | 47.5 ± 0.26 | 47.5 ± 0.18 | ** | ** | ** | 0.496 | 0.552 |
| %N | 1.5 ± 0.03 | 1.7 ± 0.03 | 1.4 ± 0.03 | 1.5 ± 0.04 | *** | *** |  |  | 0.405 |
| C:N | 29.7 ± 0.64 | 27.3 ± 0.65 | 34.1 ± 0.80 | 32 ± 0.83 | *** | ** |  |  | 0.430 |
| Collembola Abundance (individuals g dwt^-1^) | 9.2 ± 1.69 | 7.7 ± 1.33 | 8.2 ± 2.02 | 6.7 ± 2.15 |  |  |  | 0.016 | 0.095 |
| Oribatida Abundance (individuals g dwt^-1^) | 71.0 ± 7.24 | 67.7 ± 7.11 | 50.2 ± 9.75 | 36.2 ± 4.28 |  |  |  | 0.280 | 0.684 |
| Mesostigmata Abundance (individuals g dwt^-1^) | 8.4 ± 1.62 | 9.2 ± 0.92 | 9.8 ± 2.79 | 5.9 ± 1.34 |  |  |  | 0.041 | 0.177 |
| Astigmata Abundance (individuals g dwt^-1^) | 0.4 ± 0.32 | 0.0 ± 0.00 | 0.3 ± 0.1 | 0.6 ± 0.38 | na | na | na | na | na |
| Prostigmata Abundance (individuals g dwt^-1^) | 45.6 ± 5.59 | 44.8 ± 7.11 | 32.8 ± 4.95 | 28 ± 2.95 | ** |  |  |  | 0.079 |
| ‘Other’ Fauna Abundance (individuals g dwt^-1^) | 2.6 ± 0.45 | 2.1 ± 0.32 | 3 ± 0.59 | 2 ± 0.34 |  |  |  | 0.043 | 0.316 |
| Total Animal Abundance (individuals g dwt^-1^) | 137.2 ± 13.74 | 131.5 ± 13.15 | 104.3 ± 9.64 | 79.5 ± 7.05 |  |  |  | 0.215 | 0.326 |

Methods S1*.* Metabolomic signature of initial litter

A more detailed method of the procedure to analyze the metabolic signature of initial litter can be found in (Quer et al., 2022). Briefly, the extraction of the metabolites was performed on a total of 20 samples of litter (4 litter treatments × 5 replicates). A total of 10 mg of litter powder of each sample was extracted in 400 *µ*L of a MeOH/water 1:1 (v/v) mixture, followed by an ultrasound bath for 5 min at room temperature. The extracts were then centrifugated at 14000 rpm for 5 min at room temperature before analysis of the supernatants.

The metabolomic litter fingerprints were acquired on a UHPLC instrument (Dionex Ultimate 3000 RS, Thermo Scientific®, Waltham, MA, USA) equipped with a RS pump, an automatic sampler, and a thermostatically controlled column oven, and coupled to a Photodiode Array Detector (PDA), as well as an accurate quadrupole Time of Flight (qToF) mass spectrometer equipped with an Electrospray Ionisation ESI source (Impact II, Bruker Daltonics®, Billerica, MA, USA). The separation of the metabolites was carried out using a XB-C18 column (2.1 × 150 mm, 1.7 *μ*m, Phenomenex®, Torrance, CA, USA) and using a gradient elution with water and acetonitrile (MS grade, Carlo Erba®, Cornaredo, Italy), both acidified with 0.1% formic acid (Carlo Erba®, Cornaredo, Italy). The flow rate was set up at 0.6 mL min^−1^ at 35 °C.. MS detection was performed in both negative and positive ionization modes. The injection volume was 0.4 *µ*L for the acquisition in negative mode, and 1 *µ*L in positive mode.

The acquired MS-based metabolomic data were then manually calibrated using the internal calibration with the formate/acetate solution before exporting the data in *.mzXML files (centroid mode, 32 bits, zlib compression unchecked, peak picking 1-2) using MSConvert from Proteowizard. MzXML files were then corrected using the script from (Breaud et al., 2023). All the converted and modified analyses were then processed by the XCMS software (version 3.22.0; (Smith et al., 2006) under R software version (version 4.3.1; (R Core Team, 2023), following different steps to generate the final data matrix: (1) Peak picking [method="centWave", peakwidth=c(2,10), ppm=10, prefilter=c(3,100), mslevel=1, scanrange = c(100,10200)]; (2) correction of the retention times (method = obiwarp), (3) grouping (method="density", mzwid=0.01, bw=10, minfrac=0.5), (4) Fillpeaks, and then (5) report and generation of the data matrix transferred to Excel. The matrix was then filtered in three steps using an in-house R script: (1) filtering of the matrix using the Signal/Noise (S/N) ratio in order to remove the peaks observed in the blanks relatively to pooled samples (S/N = 5), (2) filtering of the matrix using the feature coefficient of variations in pooled samples in order to remove the peaks with variable intensities (threshold at 0.2), (3) filtering of the matrix according to autocorrelation between the peaks in the samples (threshold at 0.8). See Figures S8 and S9 on the following pages for example outputs of leaf metabolomic profiles.

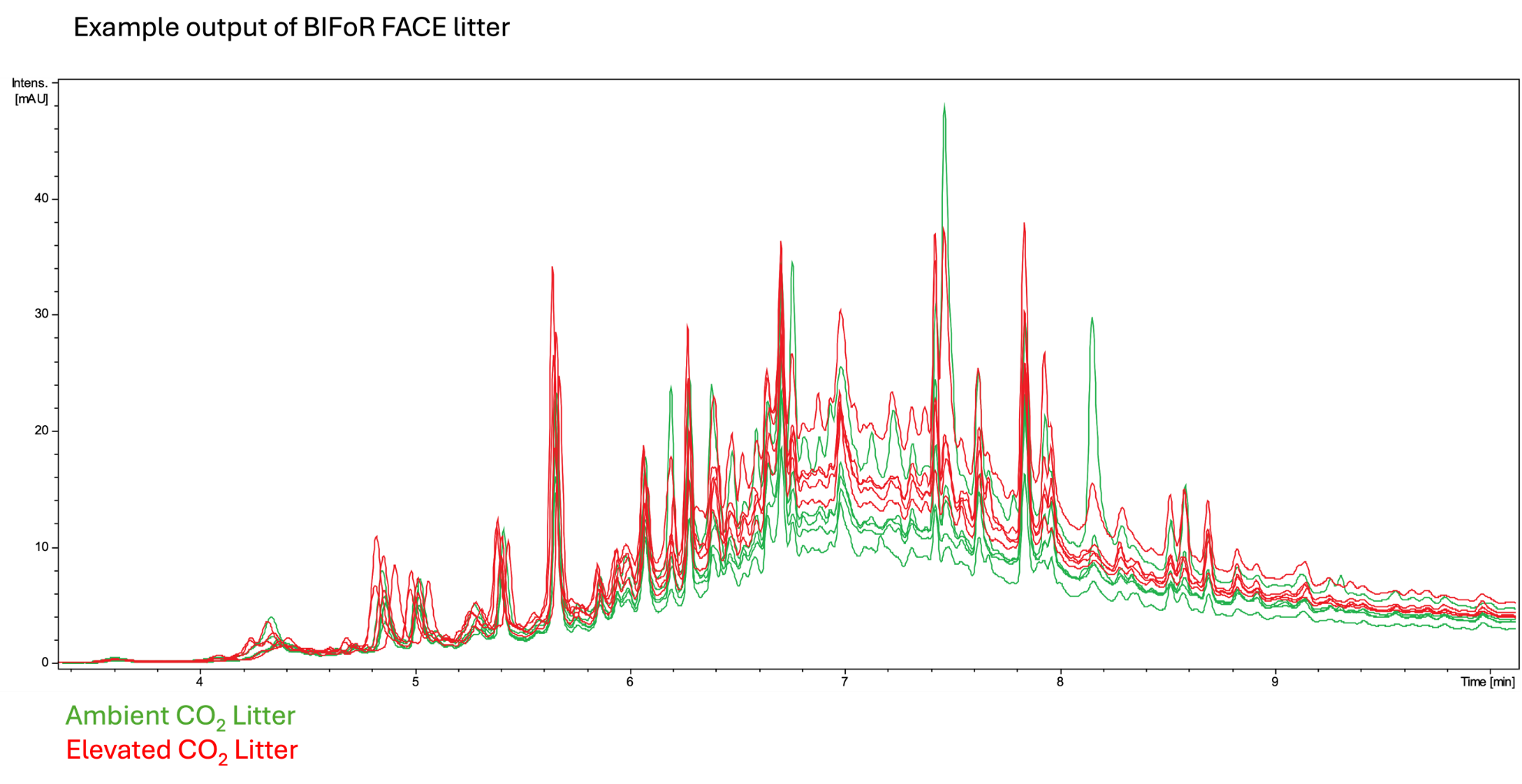

Figure. S8 Example output comparing the metabolomics profile of ambient and elevated CO_2_ initial litter from the BIFoR FACE experiment. Small differences were detected between the two litter types, which was further supported analyzing metabolomic profiles with PERMANOVA (see Figure S5).

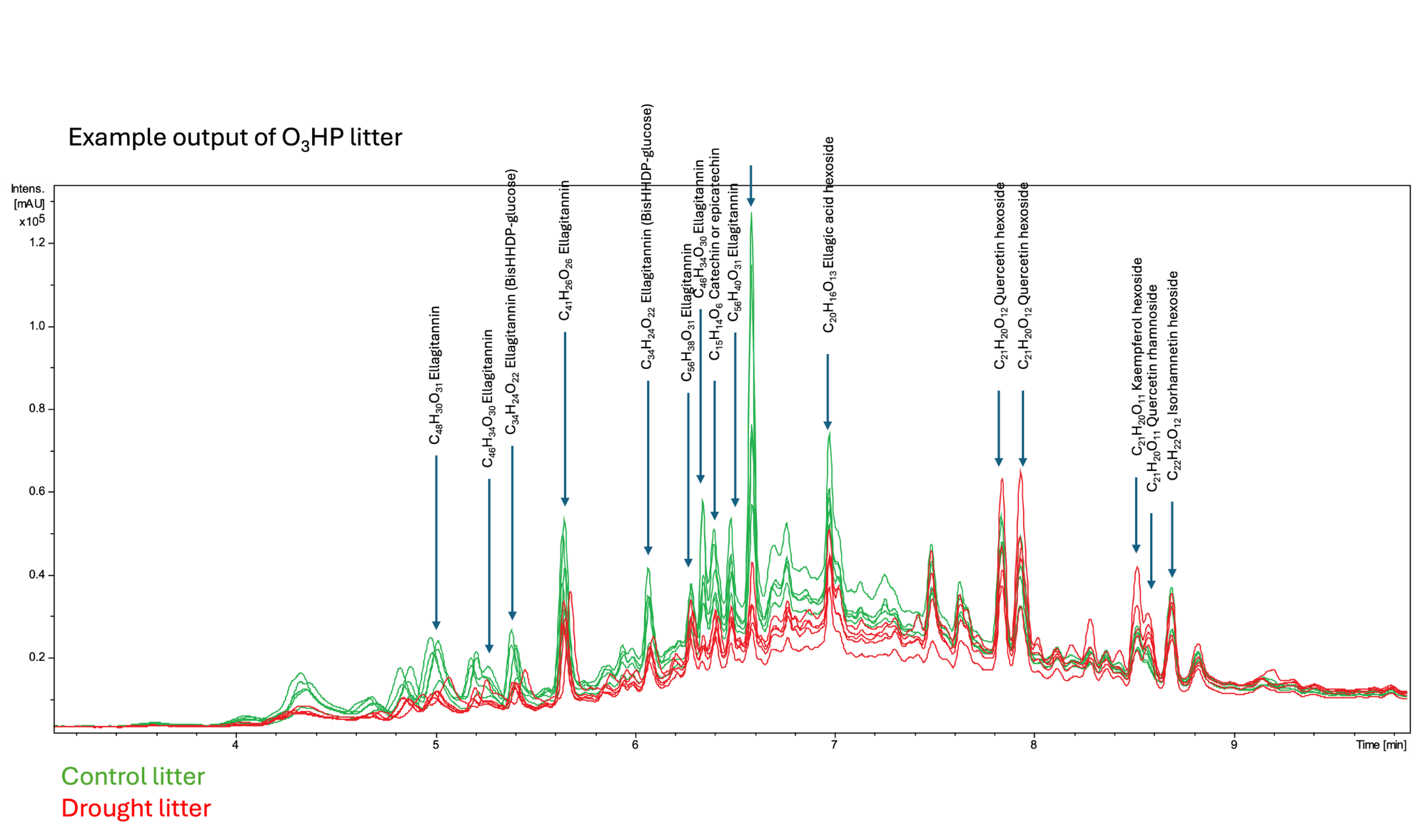

Figure. S9 Example output comparing the metabolomics profile of control and droughted initial litter from the O_3_HP experiment. Large differences were detected between the two litter types, which was further supported analyzing metabolomic profiles with PERMANOVA (see Figure S5). Specific peaks indicating metabolites (along with molecular formula) that were qualitatively different between control and droughted litter are shown.
